# Pauses in a fast-paced life: Intermittent hovering in hummingbirds

**DOI:** 10.64898/2026.04.15.716208

**Authors:** Kristiina Hurme, Ana Melisa Fernandes, Diego Andrés Benitez Duarte, Marialejandra Castro-Farías, F. Gary Stiles, Ashley Smiley, Christopher J. Clark, Alejandro Rico-Guevara

## Abstract

Hummingbirds are renowned for sustained hovering powered by continuous and rapid wingbeats. Here we describe and quantify a novel flight behavior—intermittent hovering—in which hummingbirds momentarily pause their wing motion mid-air while maintaining vertical position, with wings fully extended at the end of the upstroke. Using slow-motion footage from 86 species spanning all nine major hummingbird clades, we found that 45 species exhibited flap-pauses during sustained hovering. Phylogenetic comparative analyses revealed that hovering pauses are evolutionarily conserved and significantly associated with greater body mass and longer wings. Additionally, we found 16 species in our dataset with brightly colored underwings and they also had significantly longer wings than species with gray wings. The confluence of intermittent hovering, wing elongation, and chromatic traits lead us to hypothesize that this behavior could serve a role in visual and/or auditory communication rather than relating primarily to energy savings.

## Introduction

Hummingbirds (family Trochilidae) are unique among birds in their capacity to sustain hovering flight without relying on wind, generating lift during both the downstroke and upstroke of their extended, stiff wings (e.g., Norberg 1990; Warrick et al. 2005). This ability has made them a central model system for the study of flapping aerodynamics (Altshuler and Dudley 2002; Warrick et al. 2005; Hedrick et al. 2012). Classical intermittent flight—defined as regular alternation between flapping and non-flapping phases such as flap-bounding or flap-gliding (Tobalske 1996)—has not been considered in hovering hummingbirds, because non-flapping phases cannot generate lift and are therefore mechanically incompatible with sustained hovering. Consistent with this view, flap-pauses during stationary hovering have not previously been documented in birds (Tobalske 2010).

Hummingbirds show considerable interspecific variation in body size, wing morphology, foraging strategy, and habitat use (Sargent et al. 2021; Beltrán et al. 2022), all of which could influence whether and how a brief interruption of wing motion is physically possible or behaviorally advantageous. Additionally, several species display brightly colored underwings (Ayerbe-Quiñones 2015, Fig. 1), yet the functional significance of this coloration remains unexplored. Brief pauses during hovering could reduce motion blur and enhance the visual salience of wing coloration, potentially serving as signals in social interactions analogous to wing-based displays documented in other birds (e.g., Miller 1984; Anderson et al. 2013; Akçay and Beecher 2019).

**Figure 1.**
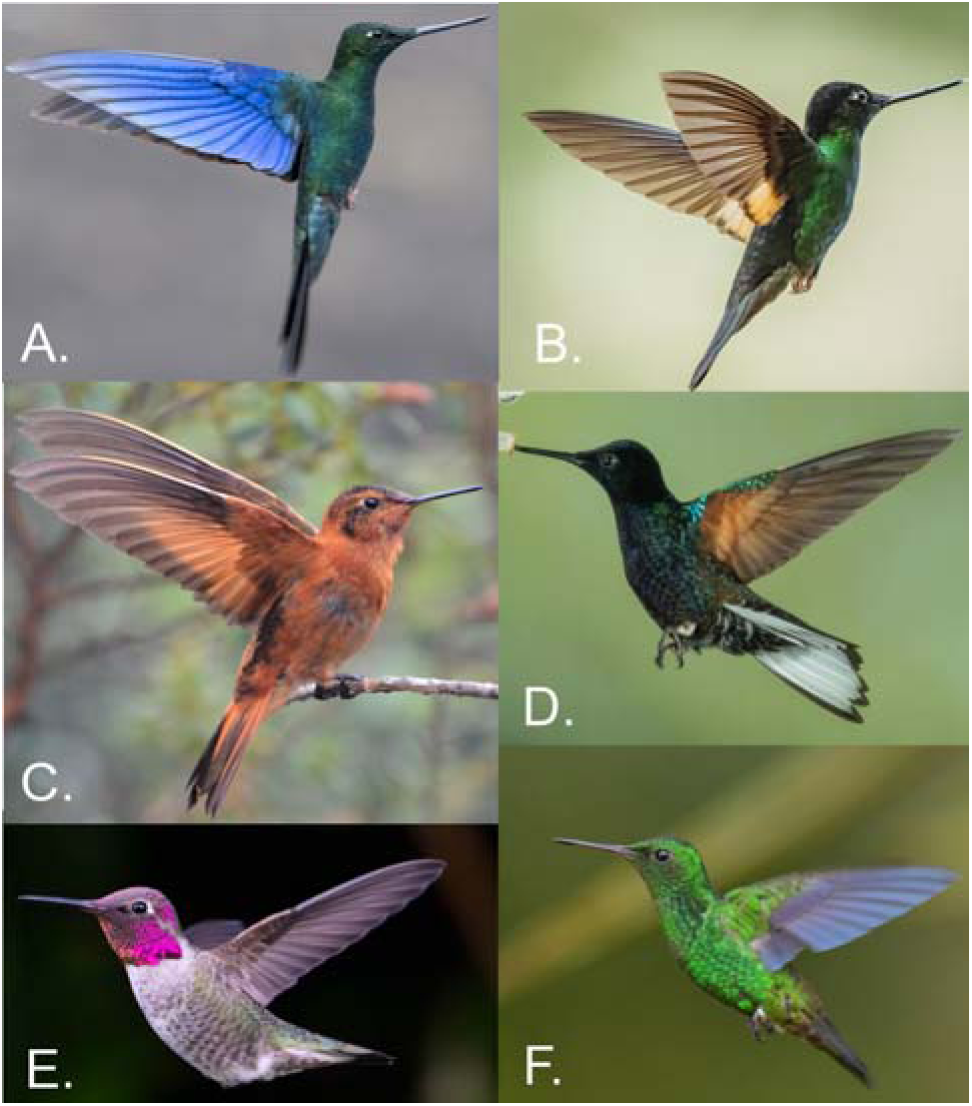
Diversity of underwing coloration in hummingbirds. Several species display shiny colors on their underwings, including blue and orange hues (e.g., A, C, D) and some translucent-colored feathers (e.g., B), while most species present gray flight feathers (e.g., E). Species and photography credits: **A.** Great sapphirewing, *Pterophanes cyanopterus* (Hank Davis), **B.** Buff-winged starfrontlet, *Coeligena lutetiae,* (Debbie Blair), **C.** Shining sunbeam, *Aglaeactis cupripennis,* (Arley Vargas), **D.** Velvet-purple coronet, *Boissonneaua jardini* (Peggy Mundy), **E.** Anna’s hummingbird, *Calypte anna,* (Alan Li), **F.** Steely-vented hummingbird*, Saucerottia saucerottei*, (Marc Faucher).

Here we describe intermittent hovering—a behavior distinct from classical intermittent flight—across 86 hummingbird species spanning all nine major clades in the family. We evaluate its phylogenetic distribution, morphological correlates, and association with underwing coloration, and discuss its potential role in visual and acoustic communication.

## Methods

We reviewed slow-motion footage of free-living hummingbirds filmed across 14 localities in 5 countries (reviewing around 180,000 frames of high-speed video at 240-500 fps), supplemented by videos from the Macaulay Library archives and other publicly available resources (e.g., social media), yielding the most phylogenetically comprehensive hovering video dataset for the family to date (Table S1). All observations were non-invasive; no animals were captured or experimentally manipulated.

We distinguish two categories of wing-motion pauses. *(1)* Hovering flap-pauses are brief arrests of wing translation and rotation occurring during sustained hovering, visible as sharply defined, immobile wings held fully extended at the end of the upstroke, with no change in altitude, gaze, or body orientation (Fig. 2A). *(2)* Transition flap-pauses occur during shifts between flight modes (e.g., deceleration of forward flight before initiating hovering or pitching forward before a descent flight) and are accompanied by visible changes in whole-body position (Fig. 2B). Pauses during forward flight—classic intermittent flight— were sporadic and excluded from species-level classification. A species was scored as pause-present (for either hovering or transition) if at least one video showed unambiguous flap-pauses, and pause-absent if none were detected after inspection of at least ten independent footage sequences exceeding 27 continuous flaps, and applied to reduce false negatives given the opportunistic nature of available footage. The final dataset comprised 86 species spanning all nine major hummingbird clades (Table S1).

**Figure 2.**
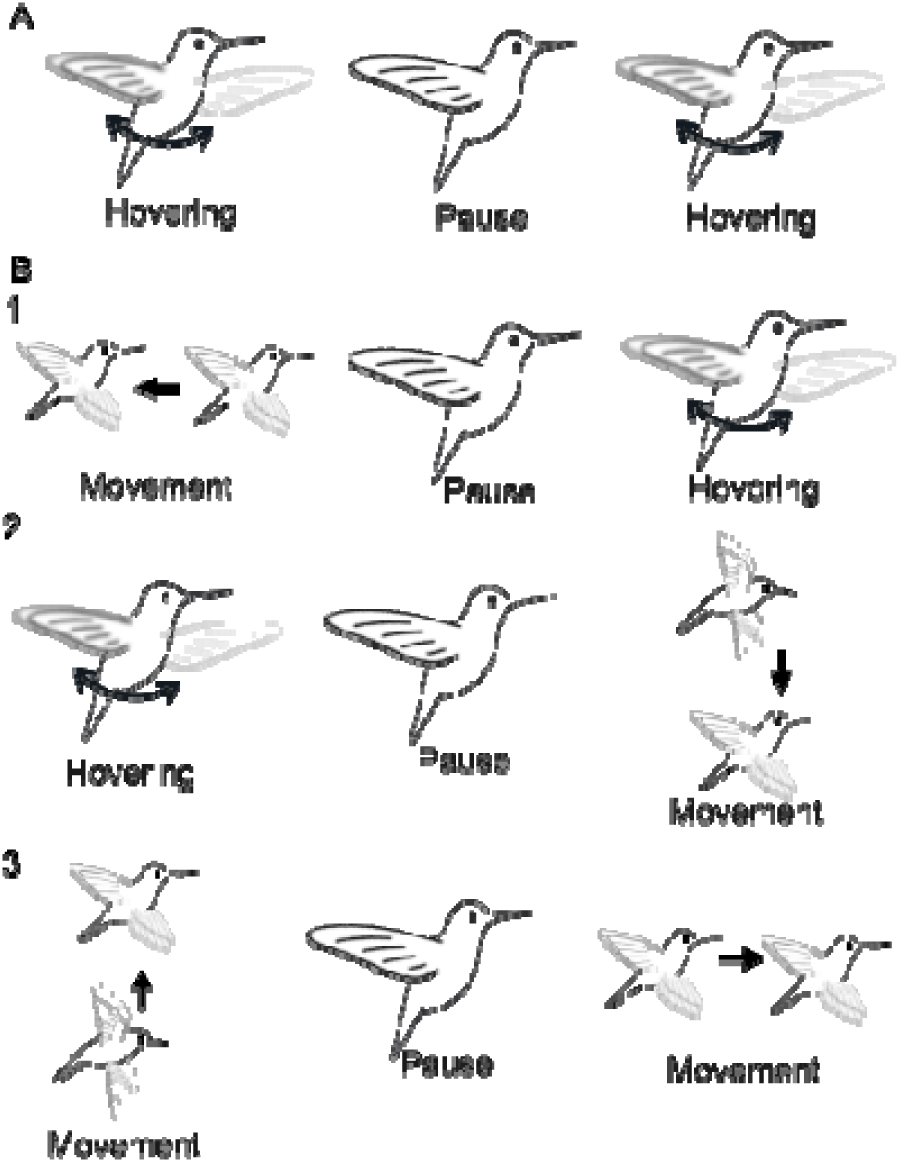
Schematic representation of intermittent hovering and flap-pause transitions in hummingbirds. **(A)** Illustration of a typical intermittent hovering sequence, showing a wingbeat cycle interrupted by a brief pause with the wings held fully extended at the end of the upstroke (Video S1). **(B)** Examples of different flap-pause transitions observed in hummingbirds. Each row depicts a distinct sequence type: transitions between hovering and directional movement in either order (rows 1 (Video S2)–2 (Video S3)), and pauses occurring between two bouts of directional movement without an intermediate hovering phase (row 3 (Video S4)). Arrows indicate the direction of wing motion and whole-body movement when present.

We scored underwing coloration (primaries and secondaries only; excluding “wingpits”) as gray or colorful from photographs in media repositories (e.g., Macaulay Library, Fig. 2), still frames from slow-motion videos, and field guides (Henderson 2010; Ayerbe-Quiñones 2015). We compiled morphological traits for the 86 study species using ranges including both sexes (Table S2). Morphological variables included body mass and length, wing length, tail length, wing area, and wing loading. We avoided including multicollinear traits (e.g., body mass, wing area, and wing loading) and evaluated them through separate phylogenetic models, rather than combining them into a single multiple regression. Morphological measurements were primarily sourced from F. G. Stiles dataset (Beltrán et al. 2022). Tail length was obtained from the database compiled by Clark (2010). Body mass data were obtained from the Colwell database (Colwell 2000), supplemented with additional values for species lacking sex-specific mass data from published sources (Montoya et al. 2018; Sheard et al. 2020; Billerman et al. 2025) and unpublished datasets (D. Plazas 2021; A. Rico-Guevara 2021; F. G. Stiles 2026). Data on pectoral type followed Zusi (2013), elevational distribution was obtained from Ayerbe-Quiñones (2015) and Rangel et al. (2015), habitat classifications from Parker et al. (1996), and foraging strategies from Rombaut et al. (2022).

All of the analyses described below were performed in R (R Core Team 2025) using the packages *ape* (Paradis and Schliep 2019), *phytools* (Revell 2024), *caper* (Orme et al. 2025), *phylolm* (Ho and Ane 2014), and *caret* (Kuhn 2008). We used the most recent time-calibrated molecular hummingbird tree (McGuire et al. 2014), pruning the phylogeny to match the species in our dataset (Paradis and Schliep 2019). Excluding the extreme size outlier—the giant hummingbird (*Patagona gigas*), we performed phylogenetic regressions to examine the evolutionary patterns underlying flap-pause occurrences, morphology, and ecology. We identified morphological predictors of hovering pauses using binary phylogenetic logistic regression (phyloglm(), MPLE method) with model selection based on AICc, evaluating univariate, additive, and interaction models. Associations between morphological variables and pause occurrence or coloration were tested with Phylogenetic Generalized Least Squares (PGLS; pgls(), incorporating phylogenetic VCV). Ancestral states were reconstructed under continuous-time Mk models (Lewis 2001), comparing Equal-Rates and All-Rates-Different models via AIC; phylogenetic signal for binary traits was quantified using Fritz & Purvis’s D statistic (phylo.d()). Correlated evolution between pauses, ecological classifications, and underwing coloration was tested using Pagel’s binary character correlation test (fitPagel()), with ecological variables binarized (Microhabitat: Open vs. Closed; Foraging: Territorial vs. Non-territorial).

## Results

### Occurrence of intermittent hovering and transition pauses

Intermittent hovering was characterized by distinct pauses in wing motion at the end of the upstroke, with wings fully extended prior to pronation; we did not observe pauses at the end of the downstroke or with wings folded against the body. Across 86 species, hovering flap-pauses were detected in 45 species (52.3%), whereas transition pauses were far more common, occurring in 78 species (90.7%). Given that only 8 species (9.3%) showed no evidence of transition pausing, they likely represent false negatives given our opportunistic sampling. Species lacking hover pauses were restricted to a few clades (Table S1, Fig. 3): bees (Mellisugini), hermits (Phaethornithini), and *Patagona*.

**Figure 3.**
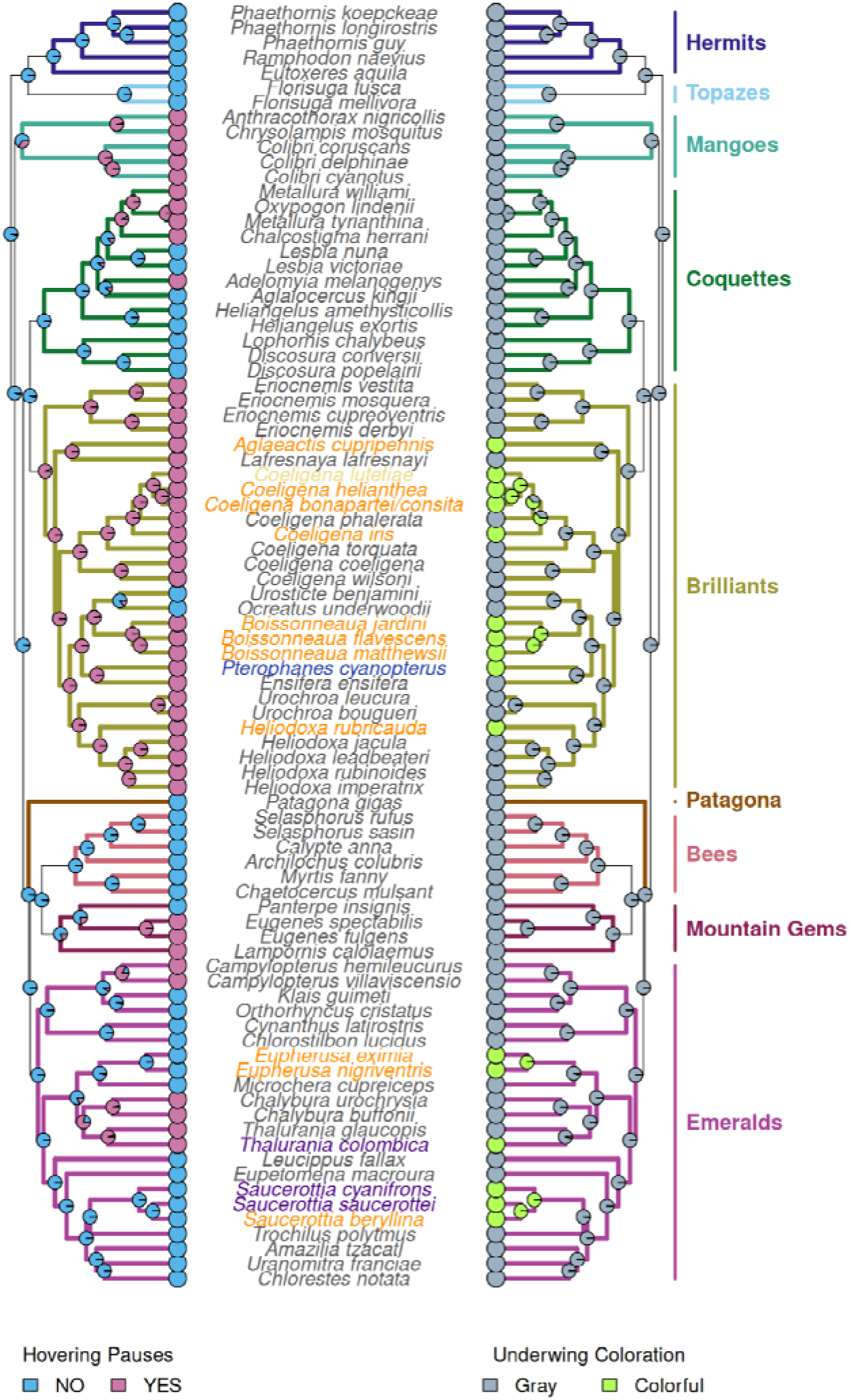
Presence of hover flap-pauses and brightly colored underwings across hummingbirds. Ancestral state reconstruction of hover pauses (left tree; pink = presence, blue = absence) and underwing coloration (right tree; gray = gray, green = colorful) across 85 hummingbird species (without *Patagona gigas*). Colored circles at the tips indicate observed states, while internal nodes show marginal ancestral state probabilities estimated via maximum likelihood. Branches are colored by clade assignment. Phylogenies were visualized using the ggtree package (Yu et al. 2017).

### Evolutionary inferences and correlates for intermittent hovering

We reconstructed the evolutionary history of hovering flap-pauses using a discrete-character maximum-likelihood model applied to the pruned phylogeny of 86 hummingbird species (Fig. 3). Ancestral state reconstructions at basal nodes indicated that the absence of hover pauses was the most likely ancestral condition. The distribution was uneven: while several lineages showed mixed reconstructions, the brilliants (Heliantheini) and mangoes (Polytminae) exhibited a distinct pattern with high support for early evolution and retention of hover pauses. In total, there appear to have been eight independent appearances of intermittent hovering along the hummingbird phylogenetic history.

Estimates of evolutionary lability under the equal-rates model indicated moderate rates of state change for hover pauses (*q* = 0.327). Phylogenetic signal analyses supported this interpretation: hover pauses exhibited strongly negative D values (*D* = -1.048, *p* < 0.001), indicating significant phylogenetic conservatism. Transition flap-pauses (Fig. S1) showed a more continuous distribution, a lower estimated transition rate (*q* = 0.171), and a D value closer to zero (*D* = 0.100, *p* = 0.01). Pagel’s tests indicated independent evolution between hover pauses and ecological variables—e.g., microhabitat (*p* = 0.677) and foraging strategy (*p* = 0.389).

Model selection based on AICc revealed univariate body mass and wing length as the best morphological predictors. Hummingbirds with longer wings and larger mass were more likely to perform hover pauses. Wing area was excluded to avoid phylogenetic collinearity with wing length (*r* = 0.917). Wing length and body mass showed significant independent effects, while their additive and sole interaction model (wing × mass) were not significant and/or performed worse. This indicates that the probability of evolving intermittent hovering is driven independently by absolute body size and wing dimensions, rather than by a synergistic biomechanical interaction.

### Underwing coloration patterns across hummingbirds

Out of 86 hummingbird species, only 16 of them (18.6%) exhibited colored underwings, with the rest having gray wings (Table S2). These colorful-winged species belonged to two clades: brilliants (Coeligenini), which contributed the largest number of orange-winged taxa—plus one blue-winged species, and emeralds (Trochilini), which contained several orange-winged species and most species with blue-hued wings. Two species, the purple-backed sunbeam (*Aglaeactis aliciae)* and the buff-winged starfrontlet (*Coeligena lutetiae*), exhibited very light or nearly white feathers in their underwings (Table S2). The ancestral state reconstruction analysis indicated that underwing coloration evolved multiple times independently (Fig. 3), each restricted to internal nodes within brilliants or emeralds. There were no species with colored wings among traplining hummingbirds.

PGLS analyses revealed that hover pauses are associated with both longer wing lengths (*p* < 0.001) and greater body mass (*p* < 0.001). Underwing coloration, by contrast, was not associated with body mass (*p* = 0.586) but showed a significant association with absolute wing length (*p* = 0.02471). This association was heavily influenced by the Great Sapphirewing (*Pterophanes cyanopterus*), an extreme outlier for wing length/area; after excluding this taxon, the relationship became only marginally significant (*p* = 0.067). Even within this reduced model, colorful-winged species had longer wings on average (adjusted mean ≈ 68.7 mm) than gray-winged species (adjusted mean ≈ 61.6 mm), though underwing coloration explained only ∼5% of the total variation in wing length (R² ≈ 0.051), with strong phylogenetic structure in the residuals (Pagel’s λ = 0.78).

Wing loading did not differ between colorful- and gray-winged species (*p* = 0.634, λ = 0.00), suggesting that the longer wings of colorful taxa are not associated with greater lift generation relative to body mass. No other assessed morphological variables differed significantly between groups (Fig. S3).

## Discussion

Here we describe a previously undocumented flight behavior in hummingbirds: flap-pausing during sustained hovering and during transitions between flight modes (Fig. 2). Hover pauses occur when the body remains stationary with only the wings moving (e.g., Video S1, S5), whereas transition pauses occur during visible shifts in body position that characterize flight mode switching (e.g., Video S2-S4).

Hover pauses show a highly probable basal state of absence, indicating that intermittent hovering arose independently on multiple occasions and became ubiquitous, or nearly so, within a few hummingbird lineages—including one apparent reversal (Fig. 3). This suggests that hover pauses represent a derived modification of flight control expressed specifically during sustained hovering. While intermittent interruptions of wing motion are known in other birds during forward flight (e.g., flap-gliding or flap-bounding), and some species capable of wind-assisted hovering (e.g., kestrels) may alternate between flapping and non-flapping phases—while gliding, pausing wing motion during sustained, unsupported hovering in still air has not previously been described. One possible explanation is that hover pauses represent responses to external perturbations such as gusts; however, their persistence in a variety of conditions without obvious perturbations—including in enclosed environments around feeders—leads us to reject this hypothesis. Instead, hover pauses likely reflect an intrinsic flight control strategy, potentially linked to fine-scale modulation of aerodynamic force production, energetic optimization, or signaling.

Hover pauses were widespread in clades of relatively large-bodied hummingbirds, such as brilliants and mangoes—with the exception of species ≤4 g—but absent in clades containing some of the smallest species, including bees and hermits (except for Costa’s hummingbird, *Calypte costae*, which performs this behavior, albeit rarely, in captivity, C.J.C. pers. obs.). Consistent with this pattern, our regression models confirm a highly significant positive association between body mass and hovering pauses (*p* < 0.01). Body mass alone, however, is not an absolute determinant: the giant hummingbird (*Patagona gigas*)—the largest species in the family—lacks intermittent hovering entirely, suggesting that other factors constrain or override the mass effect in extreme cases.

The hover pauses we document differ fundamentally from classical intermittent flight strategies. In flap-gliding and flap-bounding, non-flapping phases involve either a gliding posture or wings folded against the body during ballistic arcs (Tobalske 2010; Tobalske et al. 2011). During intermittent hovering, by contrast, pauses occur with the wings held fully extended at the end of the upstroke, with no whole-body translational movement. These kinematic differences suggest that hovering pauses are mechanistically distinct and cannot be interpreted through existing aerodynamic frameworks developed for forward flight.

Several non-mutually exclusive mechanisms could underlie hover pauses. Elastic energy storage in flight muscles and tendons has been proposed as a contributor to hummingbird hovering efficiency (Wells 1993, Chai et al. 1998, Agrawal et al. 2022), though direct measurements remain limited and its functional significance debated. Importantly, in hummingbirds the pectoralis is activated before completion of the upstroke, creating a narrow temporal separation from supracoracoideus activation; simultaneous activation of these antagonistic muscles risks tetanus and likely constrains maximum wingbeat frequency (Tobalske et al. 2010; Donovan et al. 2013). Hover pauses may therefore reflect neuromuscular constraints rather than an aerodynamic or energetic strategy—potentially achieved simply by withholding neural stimulation of the pectoralis rather than by actively controlling elastic recoil (Rayner, 1999; Tobalske 2010). Consistent with this interpretation, our comparative analyses revealed no association between hover pauses and traits indicative of elevated flight demands, and there is no physiological evidence suggesting that these pauses could reduce fatigue or improve energetic efficiency.

After accounting for phylogenetic relationships, species exhibiting hover pauses tend to have longer wings than those that do not, independently of body mass. We found no relationship with relative wing length, wing area, wing loading, or muscle composition. Together, these results suggest that absolute wing length—rather than overall size or aerodynamic scaling—is the morphological correlate of hover pauses, pointing toward functions that depend on the wing as a mechanically and/or visually salient structure.

Species with colored underwings also had longer wings when accounting for phylogeny, but the association between wing coloration and hover pauses per se was weak: many gray-winged species exhibit hover pauses (future analysis of UV coloration are warranted), and the few colorful-winged species represented in our dataset may not be enough to detect a statistical signal. Nonetheless, it is notable that most colorful-winged species do hover-pause. The exceptions—species in *Eupherusa* and *Saucerottia*—may be context-dependent: we filmed male black-bellied hummingbirds (*Eupherusa nigriventris*) performing flap-pauses with fully extended wings as part of an elaborate mating display (Video S6), suggesting that pause expression varies with behavioral context and may be underdetected in opportunistic footage. Interestingly, the oasis hummingbird (*Rhodopis vesper*, bee clade) performs similar pauses during displays but with wings closed rather than extended (Clark et al. 2013), further illustrating the behavioral diversity of wing-pause strategies across the family.

Taken together, these patterns are consistent with a signaling function for intermittent hovering. Signals are acts or structures that alter the behavior of other organisms, having evolved—or been co-opted—because of that effect, and effective because a receiver’s response has also evolved (Maynard Smith and Harper 2003). We frequently observed hover pauses in social contexts, such as around feeders where multiple individuals interact, suggesting a potential role in communication (e.g., *Boissonneaua flavescens,* video S5). Hover pauses could function to reduce the likelihood of physical confrontation by allowing individuals to assess opponent size, wing span, coloration, or flight competence. Such signaling could serve to assert dominance in competitive interactions (e.g., Falk et al 2025), deterring escalation, and avoid the high fitness costs associated with injury (e.g., by risking their bills as weapons during physical confrontation, Rico-Guevara and Araya-Salas 2015; Garzón-Agudelo et al. 2025). Such assessment signals are well documented across animal taxa, including opercular flaring in Siamese fighting fish (Portugal 2023) and parallel walking in ungulates (Jennings et al. 2003; Jennings 2012). The use of wings as signaling devices is also widespread in birds, occurring in both aggressive and courtship contexts (e.g., Miller 1984; Whitfield and Brade 1991; Anderson et al. 2013; Akçay and Beecher 2019; Mora et al. 2025). Furthermore, hovering odonates perform brief wing “standstills” that expose ornaments interpreted as signals to rivals or potential mates (Rüppell and Hilfert-Rüppell 2013).

Performing hover pauses may nonetheless carry some cost. Interrupting flapping at the end of the upstroke may require opposing elastic energy stored in stretched muscles and tendons. It remains unclear how aerodynamic force production during the pause compares to that of continuous flapping; wake recapture from the previous stroke could sustain some force generation even while the wings are momentarily arrested, hence the aerodynamic consequences of this gap need to be measured in future studies. Whether reduced wingbeat frequency is aerodynamically beneficial in some contexts remains an open question, but we note that our comparative data provide no support for an energetic benefit, and the behavior’s appearance in social contexts further weighs against an aerodynamic explanation. In stark contrast, transition pauses are widespread across hummingbirds (78 of 86 species) and likely reflect a general feature of avian flight control rather than a hummingbird-specific innovation, consistent with ancestral state reconstructions placing them near the base of the phylogeny.

In addition to their visual effects, hover pauses alter the acoustic properties of flight that produce the smooth and continuous hum (inspiring their name onomatopoeia). Continuous hovering produces sound up to approximately the sixth harmonic of wingbeat frequency (Clark and Mistick 2020), and the brief hover pauses introduce temporal gaps in sound production that were audible to human observers at close range in some species (e.g., *Boissonneaua* spp.; A.R.-G., pers. obs.). Given that hummingbirds likely possess higher temporal auditory acuity than humans in part due to their small head size (Dooling 1992; Duque et al. 2020), intermittent hovering may increase the salience of wing-generated sounds to conspecifics—consistent with the interpretation of other hummingbird wing sonations as signals (Rico-Guevara et al. 2022).

In summary, the phylogenetic distribution of intermittent hovering, its association with body mass and wing length, its occurrence in social contexts, and the lack of evidence for aerodynamic or energetic benefits are collectively more consistent with a signaling function than with an energy-saving role. Future quantification of intermittent hovering may allow testing whether hummingbirds are capable of using this behavior to communicate information about size, flight performance, competitive intent, or mate quality, potentially reducing the need for costly physical interactions. Playback experiments contrasting continuous versus intermittent hovering sounds, behavioral tracking of pause frequency across social and foraging contexts, and aerodynamic modeling of pause kinematics represent key next steps toward testing these hypotheses and evaluating the relative contributions of visual and acoustic components of this heretofore unstudied behavior.

## Data available

All source code, along with the database and phylogenetic files necessary to reproduce this analysis, is publicly hosted in this GitHub repository.

## Authors’ contributions

K.H.: conceptualization, data curation, formal analysis, investigation, methodology, project administration, validation, visualization, writing—original draft, writing—review and editing; A.M.F: conceptualization, data curation, formal analysis, investigation, methodology, project administration, validation, visualization, writing—original draft, writing—review and editing; D.A.B.D.: data curation, investigation, methodology, writing—review and editing; M.C.F.: data curation, investigation, writing—review and editing , F.G.S.: data curation, writing—review and editing; A.S.: data curation, investigation, methodology, writing— review and editing, C.J.C.: data curation, investigation, methodology, writing—review and editing, A.R.G.: conceptualization, funding acquisition, investigation, methodology, project administration, resources, supervision, validation, writing—review and editing.

All authors gave final approval for publication and agreed to be held accountable for the work performed therein.

## Acknowledgments

We thank Mateo Hernández, Edward Hurme, María José Espejo, Diana Abaunza, and Eliana Babativa for their invaluable assistance in the field and searching for videos. We are also grateful to Elsa Quicazán for helpful discussions. We thank the Macaulay Library at the Cornell Lab of Ornithology, Hank Davis, Arley Vargas, Debbie Blair, Alan Li, Marc Faucher, Peggy Mundy and Diego Calderón for providing access to audio-visual material. Alejandro Rico-Guevara thanks the Baepler Endowed Professorship, Ellen Look and Tony Cavalieri, the Washington Research Foundation, and the NSF CAREER 2440668 for support.

## Supplemental information

**Table S1.** Species with evidence of intermittent flapping pauses. For each species, we report clade, the presence or absence of hovering pauses and transition pauses, and the source of the video material.

**Table S2.** Measurements dataset of hummingbird species included in this study. For each species we report clade, muscle type, morphological traits (body mass, wing area, wing loading, wing length, tail length, and total body length), plumage coloration (presence or absence of underwing coloration and underwing color), elevational distribution (maximum and minimum elevation), foraging type, and microhabitat.

**Table S3.**
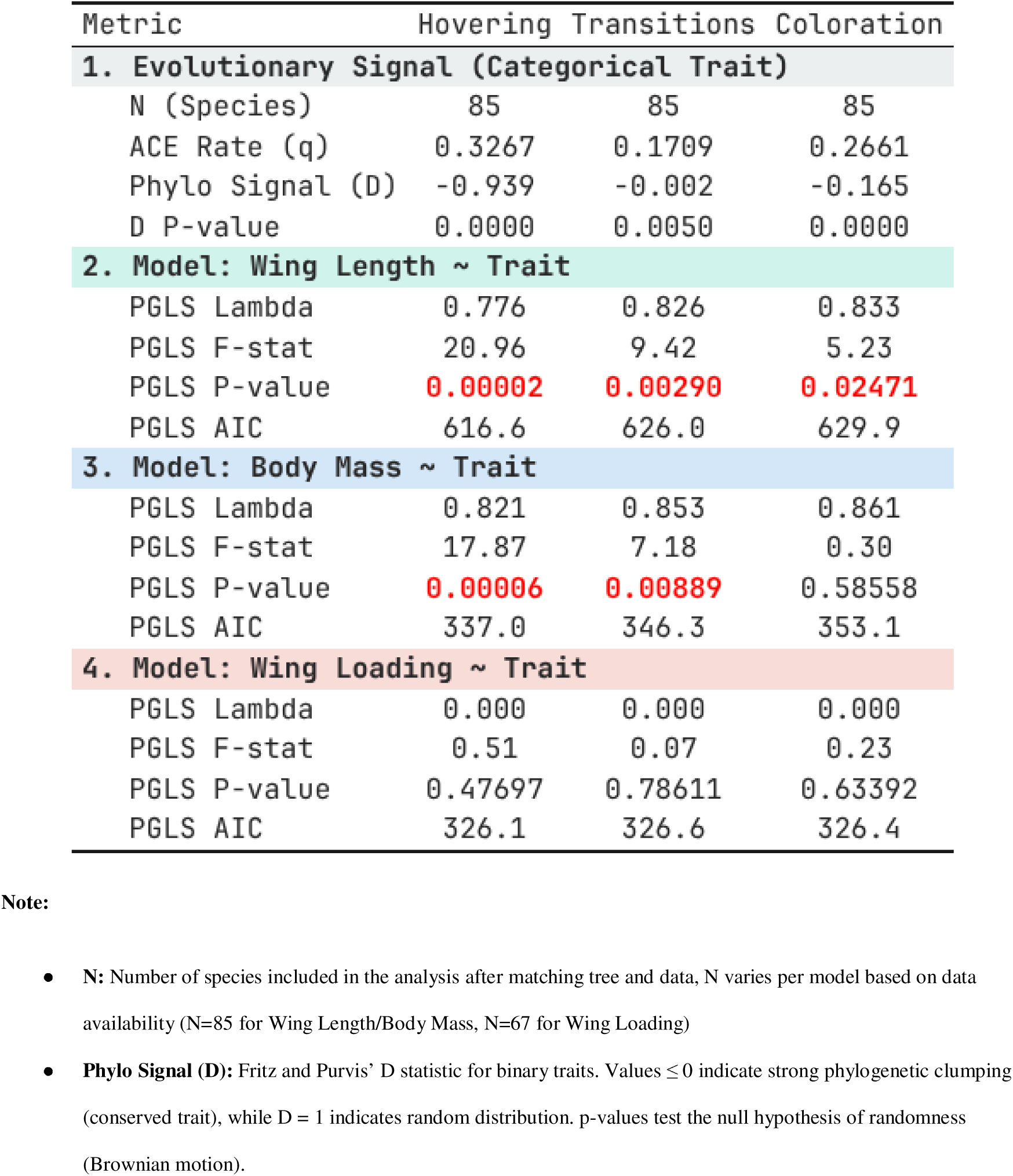
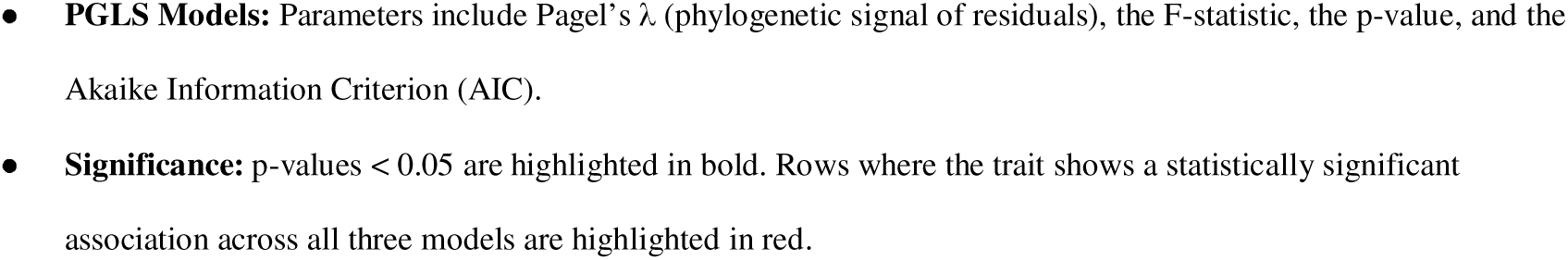
Comparative summary of evolutionary statistics and phylogenetic generalized least squares (PGLS) models testing morphological correlates of flight behavior and coloration without outliers. The table presents the estimated transition rates (q) and phylogenetic signal (D) for each binary trait, followed by PGLS model results testing the association of these traits with wing length, body mass, and the allometric relationship between wing length and body mass.

| Metric | Hovering | Transitions | Coloration |
| --- | --- | --- | --- |
| **1. Evolutionary Signal (Categorical Trait)** |  |  |  |
| N (Species) | 85 | 85 | 85 |
| ACE Rate (q) | 0.3267 | 0.1709 | 0.2661 |
| Phylo Signal (D) | -0.939 | -0.002 | -0.165 |
| D P-value | 0.0000 | 0.0050 | 0.0000 |
| **2. Model: Wing Length ~ Trait** |  |  |  |
| PGLS Lambda | 0.776 | 0.826 | 0.833 |
| PGLS F-stat | 20.96 | 9.42 | 5.23 |
| PGLS P-value | **0.00002** | **0.00290** | **0.02471** |
| PGLS AIC | 616.6 | 626.0 | 629.9 |
| **3. Model: Body Mass ~ Trait** |  |  |  |
| PGLS Lambda | 0.821 | 0.853 | 0.861 |
| PGLS F-stat | 17.87 | 7.18 | 0.30 |
| PGLS P-value | **0.00006** | **0.00889** | 0.58558 |
| PGLS AIC | 337.0 | 346.3 | 353.1 |
| **4. Model: Wing Loading ~ Trait** |  |  |  |
| PGLS Lambda | 0.000 | 0.000 | 0.000 |
| PGLS F-stat | 0.51 | 0.07 | 0.23 |
| PGLS P-value | 0.47697 | 0.78611 | 0.63392 |
| PGLS AIC | 326.1 | 326.6 | 326.4 |

**Note:**
- **N:** Number of species included in the analysis after matching tree and data, N varies per model based on data availability (N=85 for Wing Length/Body Mass, N=67 for Wing Loading) - **Phylo Signal (D):** Fritz and Purvis' D statistic for binary traits. Values $\leq 0$ indicate strong phylogenetic clumping (conserved trait), while $D = 1$ indicates random distribution. p-values test the null hypothesis of randomness (Brownian motion).
- PGLS Models:** Parameters include Pagel's $\lambda$ (phylogenetic signal of residuals), the F-statistic, the p-value, and the Akaike Information Criterion (AIC). - Significance:** p-values < 0.05 are highlighted in bold. Rows where the trait shows a statistically significant association across all three models are highlighted in red.

**Figure S1:**
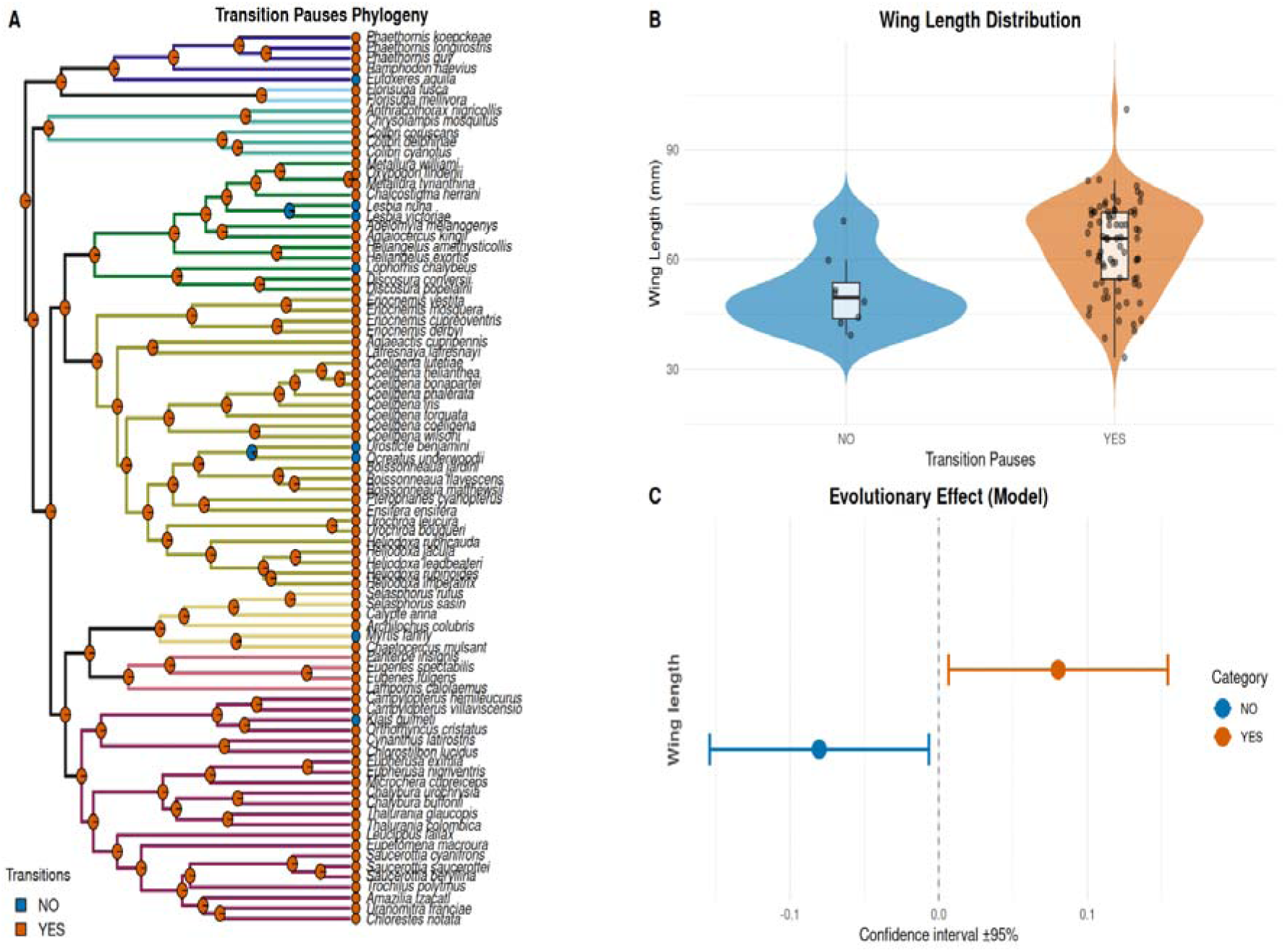
Ancestral state reconstruction and morphological correlates of transition pauses across hummingbird species. **(A)** Inferred evolutionary history of transition pauses. Internal nodes represent the marginal ancestral state probabilities for the presence (orange) and absence (blue) of the trait, estimated using a discrete-character model under maximum likelihood. The near-ubiquitous presence of this trait across the phylogeny, combined with the rare and scattered instances of its absence, suggests that its evolutionary organization is driven more by chance than by a strong phylogenetic condition. Colored circles at the tips indicate the observed states, and branches are colored by clade assignment. **(B)** Violin plots showing the distribution of wing length for species with (YES) and without (NO) transition pauses. **(C)** Phylogenetic generalized least squares (PGLS) model results testing the association between wing length and transition pauses. The plot displays the coefficient estimate ± 95% confidence intervals; intervals not crossing the dashed vertical line (zero) indicate a statistically significant association after correcting for phylogeny.

**Figure S2:**
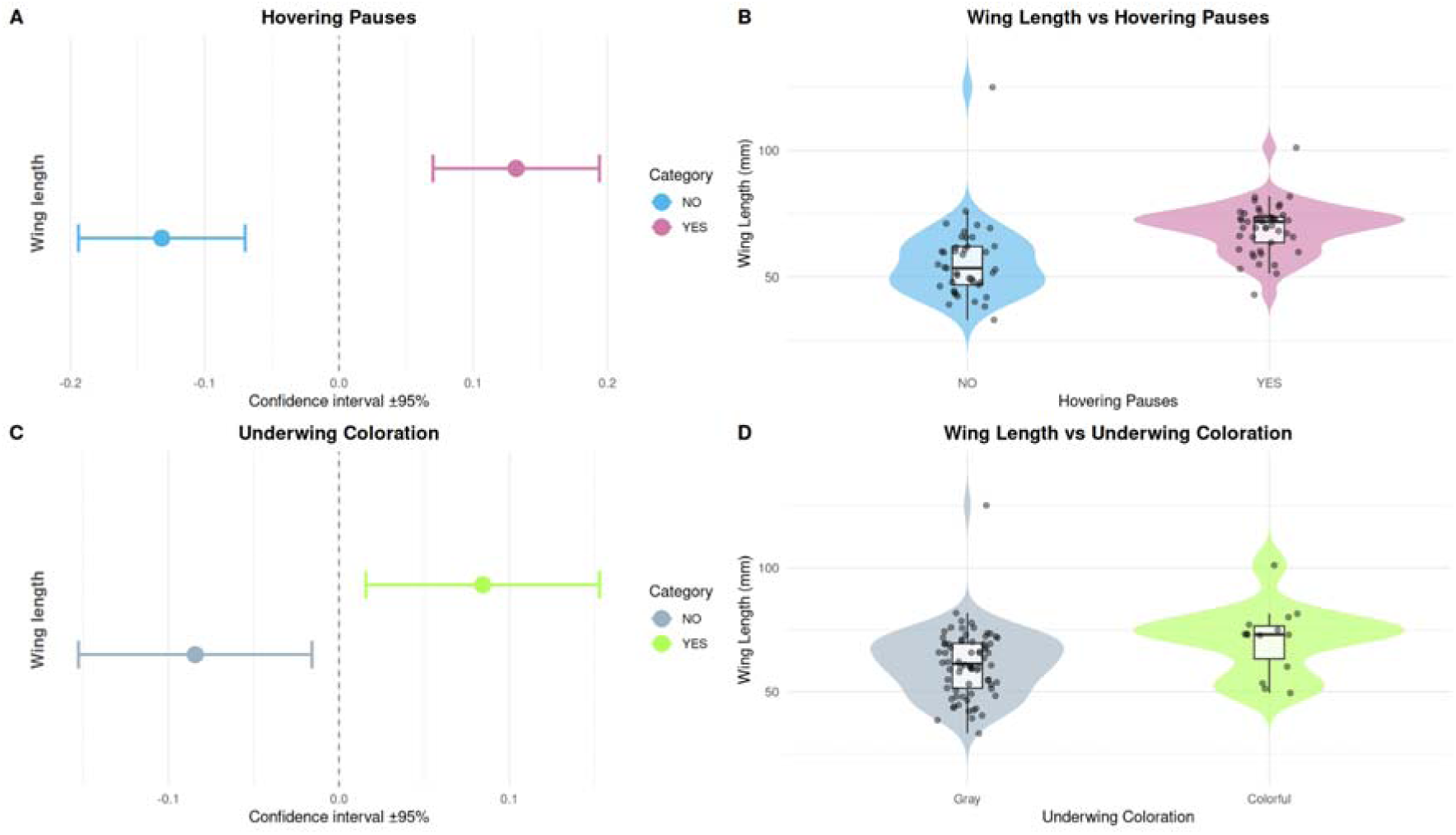
Morphological associations and phylogenetic analysis of hovering pauses and underwing coloration. **(A)** Phylogenetic logistic regression (phyloglm) model results testing the effect of wing length on the probability of hovering pause occurrence. The plot displays log-odds coefficient estimates ± 95% confidence intervals for the absence (NO, blue) and presence (YES, pink) of the behavior; divergence from the zero dashed line indicates the direction and magnitude of the evolutionary effect. **(B)** Violin plots comparing the distribution of wing length between species that do not perform hovering pauses and those that do. Each violin includes an internal boxplot (median and interquartile range) and individual data points (jitter) to visualize data dispersion. **(C)** Corresponding phylogenetic logistic regression results testing the effect of wing length on underwing coloration, showing coefficient estimates for gray (NO) and colorful (YES, green) underwings. **(D)** Violin plots comparing the distribution of wing length between species with gray and colorful underwings. Overall, the analyses demonstrate that the evolutionary acquisitions of both intermittent hovering pauses and chromatic underwing coloration are significantly associated with possessing longer wings.

**Figure S3:**
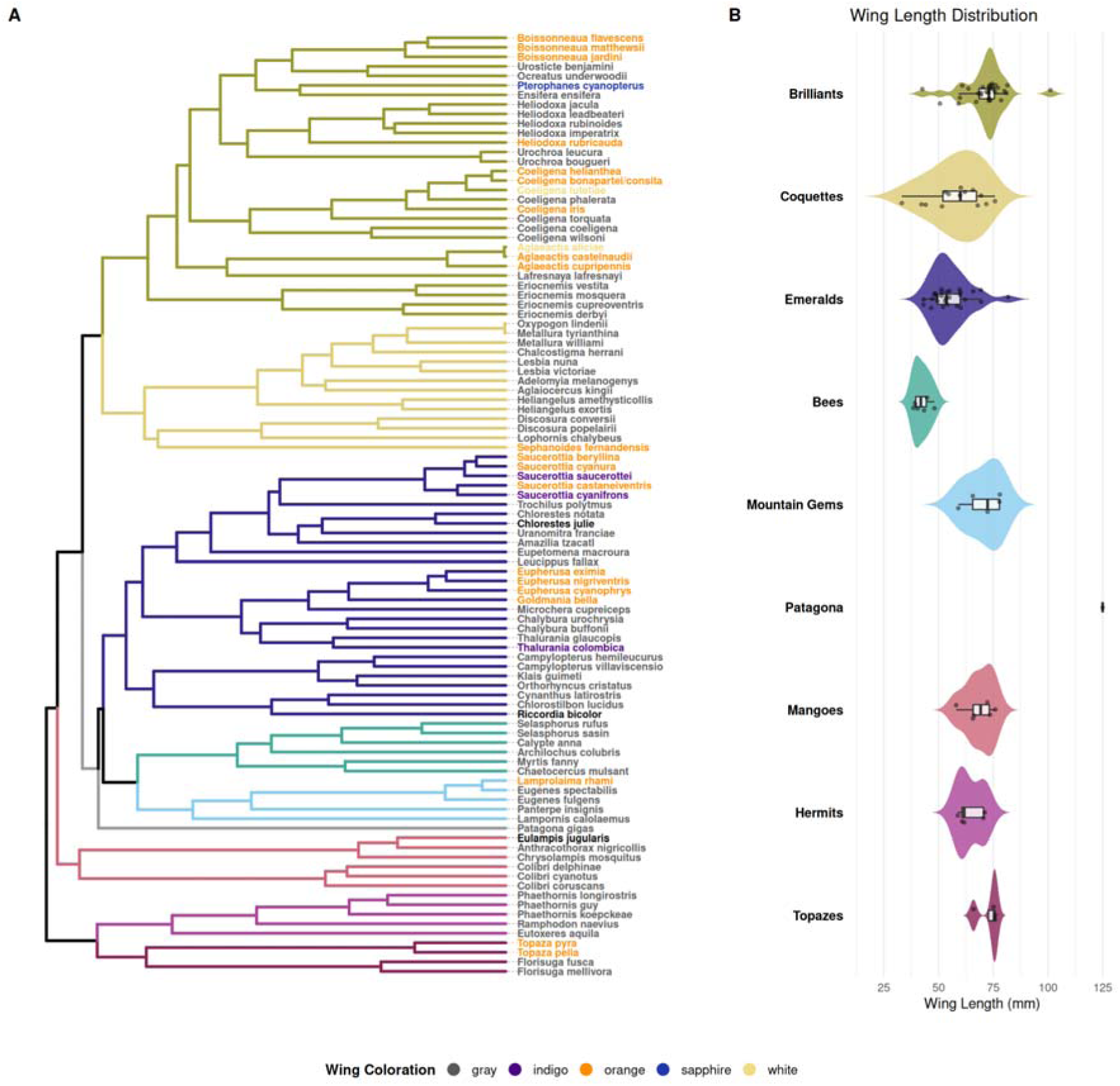
Phylogenetic distribution of underwing coloration and clade-level divergence in wing length. **(A)** Phylogenetic tree of the studied hummingbird species. Branch colors correspond to major taxonomic clades (e.g., emeralds (Trochilini), bees (Mellisugini), mountain gems (Lampornithini), etc.). Species names at the tips are colored according to their observed underwing coloration (gray, indigo, orange, sapphire, white), illustrating the taxonomic distribution of the trait across the phylogeny. **(B)** Violin plots displaying the distribution of wing length (mm) for each clade, vertically aligned with the phylogeny. Each violin represents the probability density of the data, overlaid with a boxplot indicating the median and interquartile range, along with individual data points to visualize intraspecific variation. Violin colors correspond to the clade assignments in **(A)**.

