## Supplements for "Pauses in a fast-paced life: Intermittent hovering in hummingbirds": Supplemental tables.pdf

Table 1: Table S1. Species with evidence of intermittent flapping pauses. For each species, we report clade, the presence or absence of hovering pauses and transition pauses, underwing coloration, and the source of the video material.

| Clade | Species name | Hovering pauses | Transition pauses | Underwing color | Video source |
|---|---|---|---|---|---|
| Coquettes | *Adelomyia melanogenys* | YES | YES | Gray | Javier Zurita |
| Brilliants | *Aglaeactis cupripennis* | YES | YES | Orange | Our repository |
| Coquettes | *Aglaiocercus kingii* | NO | YES | Gray | Nicolas Reusens |
| Emeralds | *Amazilia tzacatl* | NO | YES | Gray | Our repository |
| Mangoes | *Anthracothorax nigricollis* | YES | YES | Gray | Our repository |
| Bees | *Archilochus colubris* | NO | YES | Gray | Our repository |
| Brilliants | *Boissonneaua flavescens* | YES | YES | Orange | Our repository |
| Brilliants | *Boissonneaua jardini* | YES | YES | Orange | Atelopus |
| Brilliants | *Boissonneaua matthewsii* | YES | YES | Orange | ML201393591 |
| Bees | *Calypte anna* | NO | YES | Gray | Our repository |
| Emeralds | *Campylopterus hemileucurus* | YES | YES | Gray | Our repository |
| Emeralds | *Campylopterus villaviscensio* | YES | YES | Gray | Javier Zurita |
| Bees | *Chaetocercus mulsant* | NO | YES | Gray | Our repository |
| Emeralds | *Chalybura buffonii* | YES | YES | Gray | Our repository |
| Emeralds | *Chalybura urochrysia* | YES | YES | Gray | Our repository |
| Emeralds | *Chlorestes notata* | NO | YES | Gray | Joao Victor Fernandes |
| Emeralds | *Chlorostilbon lucidus* | NO | YES | Gray | ML643579014 |
| Mangoes | *Chrysolampis mosquitus* | YES | YES | Gray | Our repository |
| Brilliants | *Coeligena bonapartei* | YES | YES | Orange | Our repository |
| Brilliants | *Coeligena coeligena* | YES | YES | Gray | Norvey Esquivel |
| Brilliants | *Coeligena helianthea* | YES | YES | Orange | Our repository |
| Brilliants | *Coeligena iris* | YES | YES | Orange | ML515237401 |
| Brilliants | *Coeligena lutetiae* | YES | YES | White | Our repository |
| Brilliants | *Coeligena phalerata* | YES | YES | Gray | Our repository |
| Brilliants | *Coeligena torquata* | YES | YES | Gray | Our repository |
| Brilliants | *Coeligena wilsoni* | YES | YES | Gray | Atelopus |
| Mangoes | *Colibri coruscans* | YES | YES | Gray | Our repository |
| Mangoes | *Colibri cyanotus* | YES | YES | Gray | Our repository |
| Mangoes | *Colibri delphinae* | YES | YES | Gray | Our repository |
| Emeralds | *Cynanthus latirostris* | NO | YES | Gray | Stoil Ivanov |

| | | | | | |
|---|---|---|---|---|---|
| Coquettes | *Discosura conversii* | NO | YES | Gray | Nicolas Reusens |
| Coquettes | *Discosura popelairii* | NO | YES | Gray | NatGeo Animals, 54" |
| Brilliants | *Ensifera ensifera* | YES | YES | Gray | Our repository |
| Brilliants | *Eriocnemis cupreoventris* | YES | YES | Gray | Our repository |
| Brilliants | *Eriocnemis vestita* | YES | YES | Gray | Our repository |
| Mountain Gems | *Eugenes fulgens* | YES | YES | Gray | Our repository |
| Mountain Gems | *Eugenes spectabilis* | YES | YES | Gray | Our repository |
| Emeralds | *Eupetomena macroura* | NO | YES | Gray | Our repository |
| Emeralds | *Eupherusa eximia* | NO | YES | Orange | Our repository |
| Emeralds | *Eupherusa nigriventris* | NO | YES | Orange | Our repository |
| Hermits | *Eutoxeres aquila* | NO | NO | Gray | James Wolfe |
| Topazes | *Florisuga fusca* | NO | YES | Gray | ML614267014 |
| Topazes | *Florisuga mellivora* | NO | YES | Gray | Our repository |
| Coquettes | *Heliangelus amethysticollis* | NO | YES | Gray | Our repository |
| Brilliants | *Heliodoxa imperatrix* | YES | YES | Gray | Nicolas Reusens |
| Brilliants | *Heliodoxa jacula* | YES | YES | Gray | Our repository |
| Brilliants | *Heliodoxa leadbeateri* | YES | YES | Gray | Javier Zurita |
| Brilliants | *Heliodoxa rubinoides* | YES | YES | Gray | Atelopus |
| Brilliants | *Heliodoxa rubricauda* | YES | YES | Orange | Our repository |
| Emeralds | *Klais guimeti* | NO | NO | Gray | James Wolfe |
| Brilliants | *Lafresnaya lafresnayi* | YES | YES | Gray | Nicolas Reusens |
| Mountain Gems | *Lampornis calolaemus* | YES | YES | Gray | Our repository |
| Coquettes | *Lesbia nuna* | NO | NO | Gray | Our repository |
| Coquettes | *Lesbia victoriae* | NO | NO | Gray | Our repository |
| Emeralds | *Leucippus fallax* | NO | YES | Gray | ML631001936 |
| Coquettes | *Lophornis chalybeus* | NO | NO | Gray | Our repository |
| Coquettes | *Metallura tyrianthina* | YES | YES | Gray | Our repository |
| Emeralds | *Microchera cupreiceps* | NO | YES | Gray | Our repository |
| Bees | *Myrtis fanny* | NO | NO | Gray | Our repository |
| Brilliants | *Ocreatus underwoodii* | NO | NO | Gray | Nicolas Reusens |
| Emeralds | *Orthorhyncus cristatus* | NO | YES | Gray | ML201498771 |
| Coquettes | *Oxypogon lindenii* | YES | YES | Gray | Our repository |
| Mountain Gems | *Panterpe insignis* | NO | YES | Gray | ML567328081 |
| Patagonas | *Patagona gigas* | NO | YES | Gray | ML622519917 |
| Hermits | *Phaethornis guy* | NO | YES | Gray | Our repository |
| Hermits | *Phaethornis koepckeae* | NO | YES | Gray | Our repository |

| | | | | | |
|---|---|---|---|---|---|
| Hermits | *Phaethornis longirostris* | NO | YES | Gray | Our repository |
| Brilliants | *Pterophanes cyanopterus* | YES | YES | Sapphire | Our repository |
| Hermits | *Ramphodon naevius* | NO | YES | Gray | Our repository |
| Emeralds | *Saucerottia beryllina* | NO | YES | Orange | Panterinhaintothewild |
| Emeralds | *Saucerottia cyanifrons* | NO | YES | Indigo | Our repository |
| Emeralds | *Saucerottia saucerottei* | NO | YES | Indigo | Our repository |
| Bees | *Selasphorus rufus* | NO | YES | Gray | Our repository |
| Bees | *Selasphorus sasin* | NO | YES | Gray | ML645087570 |
| Emeralds | *Thalurania colombica* | YES | YES | Indigo | Our repository |
| Emeralds | *Thalurania glaucopis* | YES | YES | Gray | Our repository |
| Emeralds | *Trochilus polytmus* | NO | YES | Gray | Online video |
| Emeralds | *Uranomitra franciae* | NO | YES | Gray | Our repository |
| Brilliants | *Urochroa bougueri* | YES | YES | Gray | Our repository |
| Brilliants | *Urochroa leucura* | YES | YES | Gray | Javier Zurita |
| Brilliants | *Urosticte benjamini* | NO | NO | Gray | Nicolas Reusens |

Table 2: Table S2. Measurements dataset of hummingbird species included in this study. For each species we report clade, muscle type, morphological traits (body mass, wing area, wing loading, wing length, relative wing length [RELwingLength = wing length divided by the square root of body mass], tail length, and total body length), plumage coloration (presence or absence of underwing coloration, underwing color, and tail color), elevational distribution (maximum and minimum elevation), foraging type, and microhabitat.

| Species | Mt | BM | WA | WLO | WL | TL | ToL | TC | MaxA | MinA | RA | FT | MH |
|---|---|---|---|---|---|---|---|---|---|---|---|---|---|
| *A. melanogenys* | 3 | 4.9 | 10.9 | 22.4 | 53.1 | 35.4 | 88 | Brown | 3400 | 600 | 2800 | Traplining | Understory |
| *A. cupripennis* | 3 | 7.8 | 2240 | 0.2 | 81.5 | 43.6 | 114 | Brown | 4600 | 2200 | 2400 | Territorial | Mixed |
| *A. kingii* | 3 | 5 | 2092 | 0.1 | 62 | 103 | 130 | Blue | 3000 | 900 | 2100 | Opportunist | Mixed |
| *A. tzacatl* | 3 | 5 | 1006 | 0.3 | 54.9 | 32 | 98.7 | Orange | 2500 | 0 | 2500 | Territorial | Open |
| *A. nigricollis* | 2 | 7.3 | 1460 | 0.3 | 66.2 | 36.9 | 130 | Magenta | 1750 | 0 | 1750 | Opportunist | Open |
| *A. colubris* | 3 | 3.5 | 659.8 | 0.5 | 38.5 | 27.4 | 83 | Copper | 1500 | 0 | 1500 | Opportunist | Open |
| *B. flavescens* | 3 | 8.7 | 1874.9 | 0.2 | 75 | 44.9 | 118.5 | White | 3600 | 1400 | 2200 | Territorial | Mixed |
| *B. jardini* | 3 | 9 | 1994 | 0.2 | 73 | 45.4 | 120 | White | 2200 | 300 | 1900 | Territorial | Mixed |
| *B. matthewsii* | 3 | 7 | NA | NA | 72.8 | 40 | 114 | Orange | 3300 | 1200 | 2100 | Territorial | Mixed |
| *C. anna* | 4 | 4.5 | 807.2 | 0.5 | 48.1 | 31.6 | 100 | Gray | 2500 | 0 | 2500 | Territorial | Open |
| *C. hemileucurus* | 3 | 11.9 | 2481.3 | 0.2 | 81.8 | 53 | 150 | Blue | 2500 | 100 | 2400 | Opportunist | Mixed |
| *C. villaviscensio* | 3 | 7.4 | 1943 | 0.2 | 65.8 | 44.8 | 130 | Blue | 1700 | 900 | 800 | Opportunist | Understory |
| *C. mulsant* | 4 | 3.8 | 514.3 | NA | 43.4 | 24.8 | 85 | Brown | 3100 | 800 | 2300 | Opportunist | Open |
| *C. buffonii* | 3 | 6.8 | 1496.1 | 0.2 | 43.2 | 40.5 | 110 | Blue | 2000 | 0 | 2000 | Territorial | Mixed |
| *C. urochrysia* | 3 | 6.8 | 1451.4 | 0.2 | 69.4 | 39.2 | 110 | Black | 900 | 0 | 900 | Opportunist | Mixed |
| *C. notata* | 3 | 4.2 | NA | NA | 48 | 30 | 87 | Blue | 1000 | 0 | 1000 | Territorial | Open |
| *C. lucidus* | 3 | 3.5 | NA | NA | 49.2 | 27.8 | 90 | Blue | 3500 | 500 | 3000 | Opportunist | Open |
| *C. mosquitus* | 2 | 4.2 | 877.2 | 0.2 | 58 | 30.9 | 95 | Orange | 1750 | 0 | 1750 | Opportunist | Open |
| *C. bonapartei* | 3 | 6.6 | 1797.6 | 0.2 | 73.3 | 40 | 109 | Copper | 3200 | 1400 | 1800 | Traplining | Mixed |
| *C. coeligena* | 3 | 6.4 | 1696.6 | 0.2 | 70.2 | 43 | 107 | Brown | 2600 | 1000 | 1600 | Traplining | Mixed |
| *C. helianthea* | 3 | 6.6 | 1710.6 | 0.2 | 77.1 | 43.6 | 110 | Black | 3300 | 1900 | 1400 | Traplining | Mixed |
| *C. iris* | 3 | 6.9 | NA | NA | 80.1 | 43.7 | 135 | Orange | 3300 | 1500 | 1800 | Traplining | Mixed |
| *C. lutetiae* | 3 | 7.3 | 1808.6 | 0.2 | 73.4 | 36 | 140 | Black | 3600 | 2600 | 1000 | Opportunist | Understory |
| *C. phalerata* | 3 | 6.1 | 1726.3 | 0.2 | 68.2 | 41.9 | 114 | White | 3700 | 1400 | 2300 | Opportunist | Mixed |
| *C. torquata* | 3 | 7.1 | 1873 | 0.2 | 75.7 | 42.2 | 145 | Black | 3200 | 1500 | 1700 | Traplining | Understory |

| | | | | | | | | | | | | | |
|---|---|---|---|---|---|---|---|---|---|---|---|---|---|
| *C. wilsoni* | 3 | 6.7 | 1649.2 | 0.2 | 69.4 | 40.5 | 105 | Brown | 2400 | 400 | 2000 | Traplining | Mixed |
| *C. coruscans* | 2 | 8.2 | 1956.8 | 0.2 | 72.3 | 45.5 | 130 | Blue | 4500 | 400 | 4100 | Territorial | Open |
| *C. cyanotus* | 2 | 5.9 | 1467.8 | 0.2 | 65.7 | 39.1 | 97 | Blue | 3000 | 600 | 2400 | Traplining | Open |
| *C. delphinae* | 2 | 6.4 | 1647.6 | 0.2 | 73.5 | 39.4 | 120 | Brown | 2800 | 0 | 2800 | Opportunist | Mixed |
| *C. latirostris* | 3 | 3.5 | 1142.3 | NA | 51.3 | 30 | 95 | Blue | 2500 | 0 | 2500 | Opportunist | Open |
| *D. conversii* | 3 | 3 | 512.4 | 0.3 | 42.2 | 56.7 | 100 | Blue | 1400 | 60 | 1340 | Traplining | Mixed |
| *D. popelairii* | 3 | 2.5 | 424 | 0.3 | 33.2 | 20 | 100 | Blue | 1500 | 500 | 1000 | Traplining | Canopy |
| *E. ensifera* | 3 | 11.2 | 1791.5 | 0.3 | 76 | 52.9 | 135 | Black | 3500 | 1700 | 1800 | Traplining | Mixed |
| *E. cupreoventris* | 3 | 5.3 | 1264.5 | 0.2 | 60.9 | 40.6 | 98 | Blue | 3000 | 2000 | 1000 | Territorial | Open |
| *E. vestita* | 3 | 4.7 | 1204.4 | 0.2 | 59.1 | 39.5 | 90 | Blue | 4600 | 2250 | 2350 | Territorial | Open |
| *E. fulgens* | 3 | 7.9 | 2240.4 | NA | 72.4 | 44.5 | 125 | Black | 3000 | 1500 | 1500 | Opportunist | Mixed |
| *E. spectabilis* | 3 | 9.9 | 1441 | 0.3 | 77.8 | 42.3 | 130 | Black | 3000 | 1500 | 1500 | Opportunist | Open |
| *E. macroura* | 3 | 9 | 2014.8 | 0.2 | 69.3 | 80 | 160 | Blue | 1500 | 0 | 1500 | Territorial | Canopy |
| *E. eximia* | 3 | 4.3 | 856 | 0.3 | 60.1 | 33.5 | 85 | Black | 2500 | 600 | 1900 | Opportunist | Canopy |
| *E. nigriventris* | 3 | 3 | NA | NA | 49.5 | 26.3 | 80 | Black | 2100 | 600 | 1500 | Opportunist | Canopy |
| *E. aquila* | 1 | 10.6 | 2134.4 | 0.3 | 70.5 | 50 | 120 | Brown | 2100 | 300 | 1800 | Traplining | Understory |
| *F. fusca* | 2 | 8 | NA | NA | 75.9 | 42 | 110 | Black | 1400 | 0 | 1400 | Territorial | Mixed |
| *F. mellivora* | 2 | 7.4 | 1482.4 | 0.3 | 65.9 | 36.6 | 115 | White | 1600 | 0 | 1600 | Territorial | Mixed |
| *H. amethysticollis* | 3 | 6 | 1447.3 | 0.2 | 67.9 | 38.9 | 100 | Blue | 3200 | 1800 | 1400 | Territorial | Open |
| *H. imperatrix* | 3 | 8.8 | 1713 | 0.3 | 73 | 40.5 | 145 | Black | 2100 | 400 | 1700 | Territorial | Canopy |
| *H. jacula* | 3 | 9.1 | 1507.9 | 0.3 | 74.4 | 45.8 | 120 | Blue | 2000 | 300 | 1700 | Opportunist | Mixed |
| *H. leadbeateri* | 3 | 7.4 | 1543 | 0.2 | 67.2 | 45.2 | 120 | Blue | 2400 | 400 | 2000 | Territorial | Mixed |
| *H. rubinoides* | 3 | 7.8 | 1597.8 | 0.2 | 69.3 | 43.3 | 106 | Brown | 2600 | 1000 | 1600 | Opportunist | Understory |
| *H. rubricauda* | 3 | 7 | 2667 | 0.1 | 73 | 43 | 113 | Orange | 1000 | 0 | 1000 | Territorial | Mixed |
| *K. guimeti* | 3 | 2.6 | 716 | 0.2 | 48.4 | 28.4 | 78 | Blue | 1900 | 150 | 1750 | Opportunist | Mixed |
| *L. lafresnayi* | 3 | 5.3 | 1348.8 | 0.2 | 63.6 | 40.2 | 100 | White | 3700 | 1500 | 2200 | Territorial | Open |
| *L. calolaemus* | 3 | 5.4 | 710 | 0.4 | 59 | 41 | 114 | Blue | 2500 | 300 | 2200 | Opportunist | Understory |
| *L. nuna* | 3 | 4 | 932.4 | 0.2 | 51.6 | 102 | 160 | Green | 3800 | 1700 | 1500 | Opportunist | Open |
| *L. victoriae* | 3 | 5.2 | 1172.3 | 0.2 | 59.8 | 150 | 200 | Black | 4100 | 2600 | 2100 | Opportunist | Open |
| *L. fallax* | 3 | 6 | 1130 | 0.3 | 60 | 34.6 | 90 | Green | 800 | 0 | 800 | Territorial | Open |
| *L. chalybeus* | 3 | 2.5 | 932 | 0.8 | 44.1 | 29 | 75 | Brown | 1000 | 100 | 900 | Traplining | Mixed |
| *M. tyrianthina* | 3 | 3.4 | 1035.5 | 0.2 | 54.6 | 34.8 | 90 | Copper | 4200 | 600 | 3600 | Territorial | Open |
| *M. cupreiceps* | 3 | 3.3 | 1557 | 0.1 | 46.6 | 25.4 | 75 | Copper | 1500 | 300 | 1200 | Traplining | Mixed |
| *M. fanny* | 4 | 2.3 | NA | NA | 39.3 | 28 | 75 | Black | 3550 | 700 | 2850 | Traplining | Open |
| *O. underwoodii* | 3 | 3 | 544.6 | 0.3 | 42.6 | 51.1 | 120 | Blue | 3100 | 850 | 2250 | Traplining | Mixed |
| *O. cristatus* | 3 | 3 | NA | NA | 47.1 | 29.2 | 85 | Blue | 500 | 0 | 500 | Traplining | Mixed |

| | | | | | | | | | | | | | |
|---|---|---|---|---|---|---|---|---|---|---|---|---|---|
| *O. lindenii* | 3 | 4.8 | 1663.3 | 0.1 | 71.8 | 56.2 | 114 | Black | 4500 | 3600 | 900 | Territorial | Open |
| *P. insignis* | 3 | 6.2 | NA | NA | 65.6 | 43.3 | 110 | Blue | 3000 | 700 | 2300 | Territorial | Mixed |
| *P. gigas* | 3 | 18 | NA | NA | 125 | 70 | 220 | Gray | 4500 | 0 | 4500 | Territorial | Open |
| *P. guy* | 1 | 5 | 1261.5 | 0.2 | 61.7 | 50.5 | 130 | Black | 3000 | 500 | 2500 | Traplining | Understory |
| *P. koepckeae* | 1 | 5 | NA | NA | 60.7 | 65.3 | 150 | Brown | 1300 | 450 | 850 | Traplining | Understory |
| *P. longirostris* | 1 | 6 | 1370.5 | 0.2 | 58.8 | 67.8 | 120 | Black | 2500 | 0 | 2500 | Traplining | Mixed |
| *P. cyanopterus* | 3 | 10.4 | 3329.1 | 0.2 | 101.1 | 63.1 | 160 | Blue | 3700 | 2600 | 1100 | Opportunist | Mixed |
| *R. naevius* | 1 | 8.2 | NA | NA | 71.1 | 44 | 155 | Black | 500 | 0 | 500 | Traplining | Understory |
| *S. beryllina* | 3 | 4.5 | 1424.7 | NA | 53.4 | 31.8 | 95 | Orange | 3000 | 50 | 2950 | Territorial | Mixed |
| *S. cyanifrons* | 3 | 4.8 | 930 | 0.3 | 59.6 | 29.9 | 85 | Blue | 3000 | 400 | 2600 | Territorial | Open |
| *S. saucerottei* | 3 | 4.5 | 891.7 | 0.3 | 53.8 | 29.8 | 87 | Blue | 3000 | 0 | 3000 | Territorial | Open |
| *S. rufus* | 4 | 3.3 | 651.1 | 0.5 | 44.7 | 27.6 | 80 | Rufous | 3000 | 0 | 3000 | Territorial | Mixed |
| *S. sasin* | 4 | 3.1 | 470.5 | 0.5 | 40.4 | 23 | 80 | Rufous | 2650 | 0 | 2650 | Opportunist | Open |
| *T. colombica* | 3 | 4 | 891.9 | 0.2 | 51.3 | 34.8 | 95 | Blue | 2000 | 0 | 2000 | Territorial | Mixed |
| *T. glaucopis* | 3 | 4.8 | NA | NA | 54.9 | 34 | 95 | Blue | 850 | 0 | 850 | Territorial | Mixed |
| *T. polytmus* | 3 | 5.2 | NA | NA | 62.1 | 150 | 200 | Black | 2256 | 0 | 2256 | Territorial | Understory |
| *U. franciae* | 3 | 5 | 979.2 | 0.3 | 52.9 | 31.3 | 110 | Brown | 2750 | 400 | 2350 | Traplining | Mixed |
| *U. bougueri* | 3 | 11.4 | 1940 | 0.3 | 78.6 | 46.8 | 117 | White | 2800 | 500 | 2300 | Territorial | Understory |
| *U. leucura* | 3 | 7.5 | NA | NA | 73.7 | 43.5 | 135 | Green | 2800 | 1600 | 1200 | Territorial | Understory |
| *U. benjamini* | 3 | 3.9 | 863 | 0.2 | 50.6 | 36.4 | 90 | Blue | 1600 | 600 | 1000 | Territorial | Understory |
| *C. herrani* | 3 | 6.2 | 1688 | 0.2 | 69.4 | 52.4 | 120 | Black | 4000 | 2450 | 1550 | Territorial | Open |
| *E. mosquera* | 3 | 5.3 | 1735 | 0.2 | 71.9 | 55 | 115 | Green | 3600 | 1200 | 2400 | Territorial | Open |
| *E. derbyi* | 3 | 5.3 | 1221 | 0.2 | 59.8 | 32.3 | 100 | Black | 3600 | 2500 | 1100 | Territorial | Open |
| *H. exortis* | 3 | 7.5 | 1343 | 0.3 | 65.7 | 45.9 | 950 | Black | 3400 | 1400 | 2000 | Territorial | Understory |
| *M. williami* | 3 | 5 | 1338 | 0.2 | 60.1 | 40.5 | 88 | Blue | 4000 | 2100 | 1900 | Territorial | Open |
