## Supplements for "Pauses in a fast-paced life: Intermittent hovering in hummingbirds": Tables_AmNat_landscape (1).docx

**Table S1.** Species with evidence of intermittent flapping pauses. For each species, we report clade, the presence or absence of hovering pauses and transition pauses, and the source of the video material.

| Clade | Species name | Hovering pauses | Transition pauses | Video source | Link |
| --- | --- | --- | --- | --- | --- |
| Bees | Archilochus colubris | NO | YES | Our repository |  |
| Bees | Calypte anna | NO | YES | Our repository |  |
| Bees | Chaetocercus mulsant | NO | YES | Our repository |  |
| Bees | Myrtis fanny | NO | NO | Our repository |  |
| Bees | Selasphorus rufus | NO | YES | Our repository |  |
| Bees | Selasphorus sasin | NO | YES | ML645087570 | https://macaulaylibrary.org/asset/645087570 |
| Brilliants | Aglaeactis cupripennis | YES | YES | Our repository |  |
| Brilliants | Boissonneaua flavescens | YES | YES | Our repository |  |
| Brilliants | Boissonneaua jardini | YES | YES | Atelopus | https://stock.adobe.com/306032290 |
| Brilliants | Boissonneaua matthewsii | YES | YES | ML201393591 | https://macaulaylibrary.org/asset/201393591 |
| Brilliants | Coeligena bonapartei | YES | YES | Our repository |  |
| Brilliants | Coeligena coeligena | YES | YES | Norvey Esquivel | https://www.instagram.com/reel/CtpRulCATwh |
| Brilliants | Coeligena helianthea | YES | YES | Our repository |  |
| Brilliants | Coeligena iris | YES | YES | ML515237401 | https://macaulaylibrary.org/asset/515237401 |
| Brilliants | Coeligena lutetiae | YES | YES | Our repository |  |
| Brilliants | Coeligena phalerata | YES | YES | Our repository |  |
| Brilliants | Coeligena torquata | YES | YES | Our repository |  |
| Brilliants | Coeligena wilsoni | YES | YES | Atelopus | https://stock.adobe.com/306038024 |
| Brilliants | Ensifera ensifera | YES | YES | Our repository |  |
| Brilliants | Eriocnemis cupreoventris | YES | YES | Our repository |  |
| Brilliants | Eriocnemis vestita | YES | YES | Our repository |  |
| Brilliants | Heliodoxa imperatrix | YES | YES | Nicolas Reusens | https://www.instagram.com/reel/DHa83-3Mo9T/ |
| Brilliants | Heliodoxa jacula | YES | YES | Our repository |  |
| Brilliants | Heliodoxa leadbeateri | YES | YES | Javier Zurita | https://www.instagram.com/reel/C8zRs_RR_ar |
| Brilliants | Heliodoxa rubinoides | YES | YES | Atelopus | https://stock.adobe.com/306032774 |
| Brilliants | Heliodoxa rubricauda | YES | YES | Our repository |  |
| Brilliants | Lafresnaya lafresnayi | YES | YES | Nicolas Reusens | https://www.instagram.com/p/DHHiuRHim23/ |
| Brilliants | Ocreatus underwoodii | NO | NO | Nicolas Reusens | https://www.instagram.com/reel/DJkejefskx1/ |
| Brilliants | Pterophanes cyanopterus | YES | YES | Our repository |  |
| Brilliants | Urochroa bougueri | YES | YES | Our repository |  |
| Brilliants | Urochroa leucura | YES | YES | Javier Zurita | https://www.instagram.com/reel/C4VnMLbrdsZ |
| Brilliants | Urosticte benjamini | NO | NO | Nicolas Reusens | https://www.instagram.com/reel/DJbwdpUsSDG/ |
| Coquettes | Adelomyia melanogenys | YES | YES | Javier Zurita | https://www.instagram.com/reels/C4B8Qu_xSUQ/ |
| Coquettes | Aglaiocercus kingii | NO | YES | Nicolas Reusens | https://www.instagram.com/reel/DJSFDigsHuO/ |
| Coquettes | Discosura conversii | NO | YES | Nicolas Reusens | https://www.instagram.com/reel/DCvDdbZiS38 |
| Coquettes | Discosura popelairii | NO | YES | NatGeo Animals, 54" | https://youtu.be/eh4YU_Ix8pQ |
| Coquettes | Heliangelus amethysticollis | NO | YES | Our repository |  |
| Coquettes | Lesbia nuna | NO | NO | Our repository |  |
| Coquettes | Lesbia victoriae | NO | NO | Our repository |  |
| Coquettes | Lophornis chalybeus | NO | NO | Our repository |  |
| Coquettes | Metallura tyrianthina | YES | YES | Our repository |  |
| Coquettes | Oxypogon lindenii | YES | YES | Our repository |  |
| Emeralds | Amazilia tzacatl | NO | YES | Our repository |  |
| Emeralds | Campylopterus hemileucurus | YES | YES | Our repository |  |
| Emeralds | Campylopterus villaviscensio | YES | YES | Javier Zurita | https://www.instagram.com/reel/DEu8C2ZRsIh |
| Emeralds | Chalybura buffonii | YES | YES | Our repository |  |
| Emeralds | Chalybura urochrysia | YES | YES | Our repository |  |
| Emeralds | Chlorestes notata | NO | YES | Joao Victor Fernandes | https://www.instagram.com/reel/DLUqv34xBlw |
| Emeralds | Chlorostilbon lucidus | NO | YES | ML643579014 | https://macaulaylibrary.org/asset/643579014 |
| Emeralds | Cynanthus latirostris | NO | YES | Stoil Ivanov | https://www.youtube.com/watch?v=tKIwsizZZ5k |
| Emeralds | Eupetomena macroura | NO | YES | Our repository |  |
| Emeralds | Eupherusa eximia | NO | YES | Our repository |  |
| Emeralds | Eupherusa nigriventris | NO | YES | Our repository |  |
| Emeralds | Klais guimeti | NO | NO | James Wolfe | https://www.youtube.com/watch?v=Lyl9IO3kUSw |
| Emeralds | Leucippus fallax | NO | YES | ML631001936 | https://macaulaylibrary.org/asset/631001936 |
| Emeralds | Microchera cupreiceps | NO | YES | Our repository |  |
| Emeralds | Orthorhyncus cristatus | NO | YES | ML201498771 | https://macaulaylibrary.org/asset/201498771 |
| Emeralds | Saucerottia beryllina | NO | YES | Panterinha_intothewild |  |
| Emeralds | Saucerottia cyanifrons | NO | YES | Our repository |  |
| Emeralds | Saucerottia saucerottei | NO | YES | Our repository |  |
| Emeralds | Thalurania colombica | YES | YES | Our repository |  |
| Emeralds | Thalurania glaucopis | YES | YES | Our repository |  |
| Emeralds | Trochilus polytmus | NO | YES | Online video | https://www.instagram.com/reels/C10cA0cRs2T/ |
| Emeralds | Uranomitra franciae | NO | YES | Our repository |  |
| Hermits | Eutoxeres aquila | NO | NO | James Wolfe | https://youtu.be/p--0FiYdZPw |
| Hermits | Phaethornis guy | NO | YES | Our repository | https://macaulaylibrary.org/asset/622519917 |
| Hermits | Phaethornis koepckeae | NO | YES | Our repository |  |
| Hermits | Phaethornis longirostris | NO | YES | Our repository | https://macaulaylibrary.org/asset/645087570 |
| Hermits | Ramphodon naevius | NO | YES | Our repository |  |
| Mangoes | Anthracothorax nigricollis | YES | YES | Our repository |  |
| Mangoes | Chrysolampis mosquitus | YES | YES | Our repository |  |
| Mangoes | Colibri coruscans | YES | YES | Our repository |  |
| Mangoes | Colibri cyanotus | YES | YES | Our repository |  |
| Mangoes | Colibri delphinae | YES | YES | Our repository |  |
| Mountain Gems | Eugenes fulgens | YES | YES | Our repository |  |
| Mountain Gems | Eugenes spectabilis | YES | YES | Our repository |  |
| Mountain Gems | Lampornis calolaemus | YES | YES | Our repository |  |
| Mountain Gems | Panterpe insignis | NO | YES | ML567328081 | https://macaulaylibrary.org/asset/567328081 |
| Patagonas | Patagona gigas | NO | YES | ML622519917 |  |
| Topazes | Florisuga fusca | NO | YES | ML614267014 | https://macaulaylibrary.org/asset/614267014 |
| Topazes | Florisuga mellivora | NO | YES | Our repository |  |

**Table S2. Measurements dataset of hummingbird species included in this study.** For each species we report clade, muscle type, morphological traits (body mass, wing area, wing loading, wing length, tail length, and total body length), plumage coloration (presence or absence of underwing coloration, and underwing color), elevational distribution (maximum and minimum elevation), foraging type, and microhabitat.

| Species | Mass (g) | Wing area | Wing loading | wing Length | TailLength | totalLength | Color | colorful % | Wing color | MaxAlt | MinAlt | RangeAlt | Foraging type | Microhabitat |
| --- | --- | --- | --- | --- | --- | --- | --- | --- | --- | --- | --- | --- | --- | --- |
| Adelomyia melanogenys | 4.9 | 1093.9 | 0.22 | 53.1 | 35.4 | 88.0 | NO | 20 | Gray | 3400 | 600 | 2800 | Traplining | Understory |
| Aglaeactis cupripennis | 7.8 | 2240 | 0.17 | 81.5 | 43.56 | 114.0 | YES | 70 | Orange | 4600 | 2200 | 2400 | Territorial | Mixed |
| Aglaiocercus kingii | 5.0 | 2092 | 0.12 | 62.0 | 103.0 | 130.0 | NO | 20 | Gray | 3000 | 900 | 2100 | Opportunist | Mixed |
| Amazilia tzacatl | 5.0 | 1006 | 0.25 | 54.9 | 32.0 | 98.7 | NO | 20 | Gray | 2500 | 0 | 2500 | Territorial | Open |
| Anthracothorax nigricollis | 7.3 | 1460 | 0.25 | 66.17 | 36.9 | 130.0 | NO | 35 | Gray | 1750 | 0 | 1750 | Opportunist | Open |
| Archilochus colubris | 3.5 | 659.8 | 0.45 | 38.5 | 27.4 | 83.0 | NO | 20 | Gray | 1500 | 0 | 1500 | Opportunist | Open |
| Boissonneaua flavescens | 8.7 | 1874.9 | 0.23 | 75.0 | 44.9 | 118.5 | YES | 60 | Orange | 3600 | 1400 | 2200 | Territorial | Mixed |
| Boissonneaua jardini | 9.0 | 1994 | 0.23 | 73.0 | 45.4 | 120.0 | YES | 70 | Orange | 2200 | 300 | 1900 | Territorial | Mixed |
| Boissonneaua matthewsii | 7.0 |  |  | 72.8 | 40.0 | 114.0 | YES | 70 | Orange | 3300 | 1200 | 2100 | Territorial | Mixed |
| Calypte anna | 4.5 | 807.2 | 0.455 | 48.1 | 31.6 | 100.0 | NO | 20 | Gray | 2500 | 0 | 2500 | Territorial | Open |
| Campylopterus hemileucurus | 11.9 | 2481.3 | 0.24 | 81.8 | 53.0 | 150.0 | NO | 15 | Gray | 2500 | 100 | 2400 | Opportunist | Mixed |
| Campylopterus villaviscensio | 7.4 | 1943 | 0.19 | 65.8 | 44.8 | 130.0 | NO | 20 | Gray | 1700 | 900 | 800 | Opportunist | Understory |
| Chaetocercus mulsant | 3.8 | 514.3 |  | 43.41 | 24.75 | 85.0 | NO | 20 | Gray | 3100 | 800 | 2300 | Opportunist | Open |
| Chalcostigma herrani | 6.2 |  |  |  |  | 120.0 | NO | 20 | Gray | 4000 | 2450 | 1550 | Territorial | Open |
| Chalybura buffonii | 6.8 | 1496.1 | 0.23 | 43.2 | 40.5 | 110.0 | NO | 20 | Gray | 2000 | 0 | 2000 | Territorial | Mixed |
| Chalybura urochrysia | 6.8 | 1451.4 | 0.23 | 69.4 | 39.2 | 110.0 | NO | 20 | Gray | 900 | 0 | 900 | Opportunist | Mixed |
| Chlorestes notata | 4.2 |  |  | 48.0 | 30.0 | 87.0 | NO | 20 | Gray | 1000 | 0 | 1000 | Territorial | Open |
| Chlorostilbon lucidus | 3.5 |  |  | 49.2 | 27.8 | 90.0 | NO | 20 | Gray | 3500 | 500 | 3000 | Opportunist | Open |
| Chrysolampis mosquitus | 4.2 | 877.2 | 0.24 | 58.0 | 30.9 | 95.0 | NO | 20 | Gray | 1750 | 0 | 1750 | Opportunist | Open |
| Coeligena bonapartei | 6.57 | 1797.6 | 0.18 | 73.3 | 40.0 | 109.0 | YES | 60 | Orange | 3200 | 1400 | 1800 | Traplining | Mixed |
| Coeligena coeligena | 6.4 | 1696.6 | 0.19 | 70.2 | 43.0 | 107.0 | NO | 35 | Gray | 2600 | 1000 | 1600 | Traplining | Mixed |
| Coeligena helianthea | 6.6 | 1710.55 | 0.19 | 77.1 | 43.6 | 110.0 | YES | 35 | Orange | 3300 | 1900 | 1400 | Traplining | Mixed |
| Coeligena iris | 6.9 |  |  | 80.1 | 43.7 | 135.0 | YES | 70 | Orange | 3300 | 1500 | 1800 | Traplining | Mixed |
| Coeligena lutetiae | 7.3 | 1808.55 | 0.2 | 73.4 | 36.0 | 140.0 | YES | 60 | White | 3600 | 2600 | 1000 | Opportunist | Understory |
| Coeligena phalerata | 6.1 | 1726.25 | 0.18 | 68.2 | 41.9 | 114.0 | NO | 45 | Gray | 3700 | 1400 | 2300 | Opportunist | Mixed |
| Coeligena torquata | 7.1 | 1872.95 | 0.19 | 75.7 | 42.2 | 145.0 | NO | 20 | Gray | 3200 | 1500 | 1700 | Traplining | Understory |
| Coeligena wilsoni | 6.7 | 1649.15 | 0.2 | 69.4 | 40.5 | 105.0 | NO | 60 | Gray | 2400 | 400 | 2000 | Traplining | Mixed |
| Colibri coruscans | 8.2 | 1956.8 | 0.21 | 72.3 | 45.5 | 130.0 | NO | 45 | Gray | 4500 | 400 | 4100 | Territorial | Open |
| Colibri cyanotus | 5.9 | 1467.8 | 0.2 | 65.7 | 39.1 | 97.0 | NO | 45 | Gray | 3000 | 600 | 2400 | Traplining | Open |
| Colibri delphinae | 6.4 | 1647.6 | 0.19 | 73.5 | 39.4 | 120.0 | NO | 20 | Gray | 2800 | 0 | 2800 | Opportunist | Mixed |
| Cynanthus latirostris | 3.5 | 1142.27 |  | 51.3 | 30.0 | 95.0 | NO | 20 | Gray | 2500 | 0 | 2500 | Opportunist | Open |
| Discosura conversii | 3.0 | 512.4 | 0.29 | 42.2 | 56.7 | 100.0 | NO | 20 | Gray | 1400 | 60 | 1340 | Traplining | Mixed |
| Discosura popelairii | 2.5 | 424 | 0.29 | 33.2 | 20.0 | 100.0 | NO | 20 | Gray | 1500 | 500 | 1000 | Traplining | Canopy |
| Ensifera ensifera | 11.2 | 1791.5 | 0.31 | 76.0 | 52.9 | 135.0 | NO | 45 | Gray | 3500 | 1700 | 1800 | Traplining | Mixed |
| Eriocnemis cupreoventris | 5.3 | 1264.5 | 0.21 | 60.85 | 40.6 | 98.0 | NO | 20 | Gray | 3000 | 2000 | 1000 | Territorial | Open |
| Eriocnemis derbyi |  |  |  |  |  | 100.0 | NO | 20 | Gray | 3600 | 2500 | 1100 | Territorial | Open |
| Eriocnemis mosquera | 5.3 |  |  |  |  | 115.0 | NO | 20 | Gray | 3600 | 1200 | 2400 | Territorial | Open |
| Eriocnemis vestita | 4.7 | 1204.4 | 0.2 | 59.1 | 39.5 | 90.0 | NO | 20 | Gray | 4600 | 2250 | 2350 | Territorial | Open |
| Eugenes fulgens | 7.9 | 2240.37 |  | 72.4 | 44.5 | 125.0 | NO | 20 | Gray | 3000 | 1500 | 1500 | Opportunist | Mixed |
| Eugenes spectabilis | 9.9 | 1441 | 0.34 | 77.8 | 42.3 | 130.0 | NO | 20 | Gray | 3000 | 1500 | 1500 | Opportunist | Open |
| Eupetomena macroura | 9.0 | 2014.8 | 0.22 | 69.3 | 80.0 | 160.0 | NO | 20 | Gray | 1500 | 0 | 1500 | Territorial | Canopy |
| Eupherusa eximia | 4.27 | 856 | 0.25 | 60.1 | 33.5 | 85.0 | YES | 60 | Orange | 2500 | 600 | 1900 | Opportunist | Canopy |
| Eupherusa nigriventris | 3.0 |  |  | 49.5 | 26.3 | 80.0 | YES | 60 | Orange | 2100 | 600 | 1500 | Opportunist | Canopy |
| Eutoxeres aquila | 10.6 | 2134.4 | 0.25 | 70.5 | 50.0 | 120.0 | NO | 20 | Gray | 2100 | 300 | 1800 | Traplining | Understory |
| Florisuga fusca | 8.0 |  |  | 75.9 | 42.0 | 110.0 | NO | 20 | Gray | 1400 | 0 | 1400 | Territorial | Mixed |
| Florisuga mellivora | 7.4 | 1482.4 | 0.25 | 65.93 | 36.6 | 115.0 | NO | 45 | Gray | 1600 | 0 | 1600 | Territorial | Mixed |
| Heliangelus amethysticollis | 6.0 | 1447.3 | 0.21 | 67.87 | 38.9 | 100.0 | NO | 20 | Gray | 3200 | 1800 | 1400 | Territorial | Open |
| Heliangelus exortis | 7.5 |  |  |  |  | 950.0 | NO | 20 | Gray | 3400 | 1400 | 2000 | Territorial | Understory |
| Heliodoxa imperatrix | 8.8 | 1713 | 0.26 | 73.0 | 40.5 | 145.0 | NO | 20 | Gray | 2100 | 400 | 1700 | Territorial | Canopy |
| Heliodoxa jacula | 9.1 | 1507.9 | 0.3 | 74.4 | 45.8 | 120.0 | NO | 35 | Gray | 2000 | 300 | 1700 | Opportunist | Mixed |
| Heliodoxa leadbeateri | 7.4 | 1543 | 0.24 | 67.2 | 45.2 | 120.0 | NO | 20 | Gray | 2400 | 400 | 2000 | Territorial | Mixed |
| Heliodoxa rubinoides | 7.8 | 1597.8 | 0.24 | 69.3 | 43.3 | 106.0 | NO | 60 | Gray | 2600 | 1000 | 1600 | Opportunist | Understory |
| Heliodoxa rubricauda | 7.0 | 2667 | 0.13 | 73.0 | 43.03 | 113.0 | YES | 70 | Orange | 1000 | 0 | 1000 | Territorial | Mixed |
| Klais guimeti | 2.6 | 716 | 0.18 | 48.4 | 28.4 | 78.0 | NO | 20 | Gray | 1900 | 150 | 1750 | Opportunist | Mixed |
| Lafresnaya lafresnayi | 5.3 | 1348.8 | 0.2 | 63.6 | 40.2 | 100.0 | NO | 35 | Gray | 3700 | 1500 | 2200 | Territorial | Open |
| Lampornis calolaemus | 5.4 | 710 | 0.38 | 59.0 | 40.98 | 114.0 | NO | 20 | Gray | 2500 | 300 | 2200 | Opportunist | Understory |
| Lesbia nuna | 4.0 | 932.4 | 0.214500215 | 51.55 | 102.0 | 160.0 | NO | 20 | Gray | 3800 | 1700 | 1500 | Opportunist | Open |
| Lesbia victoriae | 5.2 | 1172.3 | 0.221786232 | 59.75 | 150.0 | 200.0 | NO | 20 | Gray | 4100 | 2600 | 2100 | Opportunist | Open |
| Leucippus fallax | 6.0 | 1130 | 0.265486726 | 60.0 | 34.6 | 90.0 | NO | 20 | Gray | 800 | 0 | 800 | Territorial | Open |
| Lophornis chalybeus | 2.5 | 932 ± 0.70 | 0.803 | 44.11 | 29.0 | 75.0 | NO | 20 | Gray | 1000 | 100 | 900 | Traplining | Mixed |
| Metallura tyrianthina | 3.4 | 1035.48 | 0.16 | 54.6 | 34.8 | 90.0 | NO | 20 | Gray | 4200 | 600 | 3600 | Territorial | Open |
| Metallura williami | 5.0 |  |  |  |  | 88.0 | NO | 20 | Gray | 4000 | 2100 | 1900 | Territorial | Open |
| Microchera cupreiceps | 3.3 | 1556.95 | 0.11 | 46.58 | 25.4 | 75.0 | NO | 35 | Gray | 1500 | 300 | 1200 | Traplining | Mixed |
| Myrtis fanny | 2.3 |  |  | 39.3 | 28.0 | 75.0 | NO | 20 | Gray | 3550 | 700 | 2850 | Traplining | Open |
| Ocreatus underwoodii | 3.0 | 544.6 | 0.28 | 42.6 | 51.1 | 120.0 | NO | 20 | Gray | 3100 | 850 | 2250 | Traplining | Mixed |
| Orthorhyncus cristatus | 3.0 |  |  | 47.1 | 29.2 | 85.0 | NO | 20 | Gray | 500 | 0 | 500 | Traplining | Mixed |
| Oxypogon lindenii | 4.8 | 1663.3 | 0.14 | 71.8 | 56.2 | 114.0 | NO | 20 | Gray | 4500 | 3600 | 900 | Territorial | Open |
| Panterpe insignis | 6.2 |  |  | 65.6 | 43.3 | 110.0 | NO | 20 | Gray | 3000 | 700 | 2300 | Territorial | Mixed |
| Patagona gigas | 18.0 |  |  | 125.0 | 70.0 | 220.0 | NO | 20 | Gray | 4500 | 0 | 4500 | Territorial | Open |
| Phaethornis guy | 5.0 | 1261.5 | 0.198176774 | 61.7 | 50.5 | 130.0 | NO | 20 | Gray | 3000 | 500 | 2500 | Traplining | Understory |
| Phaethornis koepckeae | 5.0 |  |  | 60.7 | 65.3 | 150.0 | NO | 20 | Gray | 1300 | 450 | 850 | Traplining | Understory |
| Phaethornis longirostris | 6.0 | 1370.45 | 0.22 | 58.75 | 67.8 | 120.0 | NO | 10 | Gray | 2500 | 0 | 2500 | Traplining | Mixed |
| Pterophanes cyanopterus | 10.4 | 3329.1 | 0.16 | 101.1 | 63.1 | 160.0 | YES | 95 | Sapphire | 3700 | 2600 | 1100 | Opportunist | Mixed |
| Ramphodon naevius | 8.2 |  |  | 71.1 | 44.0 | 155.0 | NO | 20 | Gray | 500 | 0 | 500 | Traplining | Understory |
| Saucerottia beryllina | 4.5 | 1424.7 |  | 53.4 | 31.8 | 95.0 | YES | 70 | Orange | 3000 | 50 | 2950 | Territorial | Mixed |
| Saucerottia cyanifrons | 4.8 | 930 | 0.26 | 59.58 | 29.9 | 85.0 | NO | 20 | Indigo | 3000 | 400 | 2600 | Territorial | Open |
| Saucerottia saucerottei | 4.5 | 891.7 | 0.25 | 53.78 | 29.8 | 87.0 | NO | 60 | Indigo | 3000 | 0 | 3000 | Territorial | Open |
| Selasphorus rufus | 3.27 | 651.1 | 0.477 | 44.72 | 27.6 | 80.0 | NO | 35 | Gray | 3000 | 0 | 3000 | Territorial | Mixed |
| Selasphorus sasin | 3.1 | 470.5 | 0.468 | 40.4 | 23.0 | 80.0 | NO | 35 | Gray | 2650 | 0 | 2650 | Opportunist | Open |
| Thalurania colombica | 4.0 | 891.9 | 0.22 | 51.3 | 34.8 | 95.0 | YES | 60 | Indigo | 2000 | 0 | 2000 | Territorial | Mixed |
| Thalurania glaucopis | 4.8 |  |  | 54.9 | 34.0 | 95.0 | NO | 20 | Gray | 850 | 0 | 850 | Territorial | Mixed |
| Trochilus polytmus | 5.19 |  |  | 62.1 | 150.0 | 200.0 | NO | 20 | Gray | 2256 | 0 | 2256 | Territorial | Understory |
| Uranomitra franciae | 5.0 | 979.2 | 0.26 | 52.9 | 31.3 | 110.0 | NO | 20 | Gray | 2750 | 400 | 2350 | Traplining | Mixed |
| Urochroa bougueri | 11.4 | 1940 | 0.29 | 78.6 | 46.8 | 117.0 | NO | 20 | Gray | 2800 | 500 | 2300 | Territorial | Understory |
| Urochroa leucura | 7.5 |  |  | 73.7 | 43.5 | 135.0 | NO | 20 | Gray | 2800 | 1600 | 1200 | Territorial | Understory |
| Urosticte benjamini | 3.9 | 863 | 0.23 | 50.6 | 36.4 | 90.0 | NO | 20 | Gray | 1600 | 600 | 1000 | Territorial | Understory |
